# Assembly of recombinant tau into filaments identical to those of Alzheimer’s disease and chronic traumatic encephalopathy

**DOI:** 10.1101/2021.12.16.472950

**Authors:** Sofia Lövestam, Fujiet Adrian Koh, Bart van Knippenberg, Abhay Kotecha, Alexey G. Murzin, Michel Goedert, Sjors H.W. Scheres

## Abstract

Abundant filamentous inclusions of tau are characteristic of more than 20 neurodegenerative diseases that are collectively termed tauopathies. Electron cryo-microscopy (cryo-EM) structures of tau amyloid filaments from human brain revealed that distinct tau folds characterise many different diseases. A lack of laboratory-based model systems to generate these structures has hampered efforts to uncover the molecular mechanisms that underlie tauopathies. Here, we report *in vitro* assembly conditions with recombinant tau that replicate the structures of filaments from both Alzheimer’s disease (AD) and chronic traumatic encephalopathy (CTE), as determined by cryo-EM. Our results suggest that post-translational modifications of tau modulate filament assembly, and that previously observed additional densities in AD and CTE filaments may arise from the presence of inorganic salts, like phosphates and sodium chloride. *In vitro* assembly of tau into disease-relevant filaments will facilitate studies to determine their roles in different diseases, as well as the development of compounds that specifically bind to these structures or prevent their formation.

## Introduction

Six tau isoforms that range from 352 to 441 amino acids in length are expressed in adult human brain from a single gene by alternative mRNA splicing (Goedert *et al*., 1989). The tau sequence can be separated into an N-terminal projection domain (residues 1-150), a proline-rich region (residues 151-243), the microtubule-binding repeat region (residues 244-368) and a C-terminal domain (residues 369-441). Isoforms differ by the presence or absence of one or two inserts of 29 amino acids each, in the N-terminal region (0N, 1N or 2N), and the presence or absence of the second of four microtubule-binding repeats of 31 or 32 amino acids each (resulting in three-repeat, 3R, and four-repeat, 4R, isoforms). The second N-terminal insert is only expressed together with the first. Tau filaments can differ in isoform composition in disease (Goedert, Eisenberg and Crowther, 2017). Thus, a mixture of all six isoforms is present in the tau filaments of Alzheimer’s disease (AD), chronic traumatic encephalopathy (CTE) and other diseases; in Pick’s disease (PiD), filaments are composed of only 3R tau isoforms; and in progressive supranuclear palsy (PSP), corticobasal degeneration (CBD), globular glial tauopathy (GGT), argyrophilic grain disease (AGD) and other tauopathies, filaments are composed of only 4R tau isoforms.

Tau filaments consist of a structured core made mostly of the repeat domain, with less structured N- and C-terminal regions forming the fuzzy coat (Goedert *et al*., 1988; Wischik *et al*., 1988b). Previously, we and others used electron cryo-microscopy (cryo-EM) to determine the atomic structures of the cores of tau filaments extracted from the brains of individuals with a number of neurodegenerative diseases (Fitzpatrick *et al*., 2017; Falcon, Zhang, Murzin, *et al*., 2018; Falcon *et al*., 2019, 2019; Arakhamia *et al*., 2020; Zhang *et al*., 2020; Shi *et al*., 2021). Whereas tau filaments from different individuals with a given disease share the same structures, different diseases tend to be characterised by distinct tau folds. Identical protofilaments can arrange in different ways to give rise to distinct filaments (ultrastructural polymorphs). In AD, two protofilaments can combine in a symmetrical way to form paired helical filaments (PHFs), or in an asymmetrical manner to form straight filaments (SFs). The extent of the ordered cores of the protofilament folds explains the isoform composition of tau filaments. Alzheimer and CTE folds consist almost entirely of R3 and R4; the Pick fold comprises most of R1, as well as all of R3 and R4; and the CBD, PSP, GGT and AGD folds comprise the whole of R2, R3 and R4. All known tau folds also contain residues 369-379/380 from the C-terminal domain. Based on these findings, we proposed a hierarchical classification of tauopathies, which complements clinical and neuropathological characterisation, and allows identification of new disease entities (Shi *et al*., 2021). Nevertheless, it remains unclear what roles distinct tau folds play in different diseases and what factors determine structural specificity.

In human diseases, full-length tau assembles into filaments intracellularly (Goedert, Eisenberg and Crowther, 2017). Proteolytic cleavage of the fuzzy coat is characteristic of extracellular ghost tangles (Endoh *et al*., 1993). Yet, full-length recombinant tau is highly soluble and *in vitro* assembly into filaments requires the addition of anionic co-factors, such as sulphated glycosaminoglycans, RNA, fatty acids and polyglutamate (Goedert *et al*., 1996; Kampers *et al*., 1996; Pérez *et al*., 1996; Wilson and Binder, 1997; Friedhoff *et al*., 1998) or the use of harsh shaking conditions with small beads and sodium azide (Chakraborty *et al*., 2021). Whether the structures of filaments assembled *in vitro* resemble those observed in disease requires verification by structure determination using solid-state NMR or cryo-EM. We previously showed that the addition of heparin to full-length 3R or 4R recombinant tau expressed in *E*.*coli* leads to the formation of polymorphic filaments, with structures that are unlike those in disease (Zhang *et al*., 2019). Below, we show that the addition of RNA or phosphoserine to full-length recombinant 4R tau also leads to filament structures that are different from those observed in disease.

The mechanisms of templated seeding, where filaments provide a template for the incorporation of tau monomers, thus resulting in filament growth, underlies the hypothesis of prion-like spreading of tau pathology. It is therefore possible that *in vitro* assembly of disease-relevant structures may also require the presence of a seed. Protein misfolding cyclic amplification (PMCA) and real-time quaking-induced conversion (RT-QuIC) are commonly used (Soto, Saborio and Anderes, 2002; Saijo *et al*., 2020). These techniques use protein aggregates from brain as a template to seed the assembly of filaments from recombinant proteins. However, they may not always amplify the predominant filamentous assemblies Therefore, structural verification of seeds and seeded aggregates is required. For α-synuclein and immunoglobulin light-chain, cryo-EM has shown that using similar amplification methods, the structures of seeded filaments were not the same as those of the seeds (Burger *et al*., 2021; Lövestam *et al*., 2021; Radamaker *et al*., 2021).

Full-length human tau is highly soluble, but truncated proteins encompassing the repeat region readily assemble into filaments that resemble AD PHFs by negative staining. A fragment consisting of residues 250-378, excluding R2, formed filaments when using the hanging drop approach, a method commonly used in protein crystallography (Crowther *et al*., 1992). The same was true of K11 and K12 constructs (tau residues 244-394 for K11, and the same sequences, excluding R2, for K12) (Wille *et al*., 1992). Moreover, proteins comprising the ordered cores of tau filaments from AD [residues 306-378], PiD [residues 254-378, but excluding R2] and CBD [residues 274-380] have been reported to assemble into filaments (Carlomagno *et al*., 2021). It remains to be established if the structures of these filaments resembled those from diseased brain. Multiple studies have also used constructs K18 and K19, which comprise the microtubule-binding repeats and 4 amino acids after the repeats (residues 244-372 for K18, and the same sequences, excluding R2, for K19); they spontaneously assemble into filaments (Gustke *et al*., 1994; Mukrasch *et al*., 2005; von Bergen *et al*., 2006; Li *et al*., 2009; Yu *et al*., 2012; Shammas *et al*., 2015) and are seeding-active by RT-QuIC (Saijo *et al*., 2020). However, since all known tau structures from human brain include residues beyond 372, tau filament structures of K18 and K19 cannot be the same as those in disease. Another fragment that has been used for *in vitro* assembly studies is dGAE, which comprises residues 297-391 of 4R tau, and was identified as the proteolytically stable core of PHFs from tangle fragments of AD (Wischik *et al*., 1988a). It assembles into filaments with similar morphologies to AD PHFs by negative staining (Novak, Kabat and Wischik, 1993; Al-Hilaly *et al*., 2017, 2020; Lutter *et al*., 2021).

Here, we report conditions that lead to the formation of AD PHFs from purified tau(297-391) expressed in *E*.*coli*. We show that the same construct can also be used to form type II filaments of CTE, and demonstrate how different cations affect the differences between both structures. In addition, we describe to what extent the tau(297-391) fragment can be extended, as well as shortened, while still forming PHFs. We report 76 cryo-EM structures, including those of 27 previously unobserved filaments. **Table 1** gives an overview of assembly conditions (numbered) and filament types (indicated with letters). **Figure 1** and **Figure 1 - figure supplement 1** describe the cryo-EM structures that were determined. Our results illustrate how high-throughput cryo-EM structure determination can guide the quest for understanding the molecular mechanisms of amyloid filament formation.

**Table 1:** *In vitro* assembly conditions for all filament types.

| Filament types | Construct (residues) | Buffer | Shaking (rpm) | Time (hrs) |
| --- | --- | --- | --- | --- |
| 1a | 297-391 | 10 mM PB 10 mM DTT pH 7.4 | 700 | 48 |
| 2a-d | 297-391 | 10 mM PB 10 mM DTT pH 7.4 | 200 | 48 |
| 3a | 297-391 | 10 mM PB 10 mM DTT pH 7.4<br>0.1 µg /ml dextran sulfate | 200 | 48 |
| 4a | 297-391 | 10 mM PB 10 mM DTT pH 7.4 200 mM MgCl<sub>2</sub> | 200 | 48 |
| 5a | 297-391 | 10 mM PB 10 mM DTT pH 7.4 20 mM CaCl<sub>2</sub> | 200 | 48 |
| 6a-c | 266/297-391* | 10 mM PB 10 mM DTT pH 7.4 | 200 | 48 |
| 7a-b | 266-273 -391 | 10 mM PB 10 mM DTT pH 7.4 200 mM MgCl<sub>2</sub> | 200 | 48 |
| 8a-b | 266/297-391 | 10 mM PB pH 7.4 10 mM DTT 200 mM NaCl | 200 | 48 |
| 9a-b | 266/297-391 | 10 mM PB pH 7.4 10 mM DTT 200 mM LiCl | 200 | 48 |
| 10a-b | 266/297-391 | 10 mM PB pH 7.4 10 mM DTT 200 mM KCl | 200 | 48 |
| 11a | 266/297-391 | 10 mM PB pH 7.4 10 mM DTT 100 µM ZnCl<sub>2</sub> | 200 | 48 |
| 12a | 266/297-391 | 10 mM PB pH 7.4 10 mM DTT 200 µM CuCl<sub>2</sub> | 200 | 48 |
| 13a | 266/297-391 | 10 mM PB pH 7.4 10 mM DTT 20 mM MgCl<sub>2</sub> 50 mM KCl 50 mM NaCl | 200 | 48 |
| 14a-b | 266/297-391 | 10 mM PB pH 7.4 10 mM DTT 20 mM MgCl<sub>2</sub> 100 mM NaCl | 200 | 48 |
| 15a-d | 266/297-391 | 10 mM PB pH 7.4 10 mM DTT 10 mM MgSO<sub>4</sub> 100 mM NaCl | 200 | 48 |
| 16a-b | 266/297-391 | 10 mM PB pH 7.4 10 mM DTT 10 mM NaHCO<sub>3</sub> 100 mM NaCl | 200 | 48 |
| 17a-c | 266/297-391 | 10 mM PB pH 7.4 10 mM DTT 500 mM NaCl | 200 | 48 |
| 18a | 244-391 | 50 mM PB pH 7.4 10 mM DTT 20 mM MgCl<sub>2</sub> | 200 | 76 |
| 19a | 244-391 | 50 mM PB pH 7.4 10 mM DTT 200 mM NaCl | 200 | 76 |
| 20a | 244-391 | 10 mM PB 10 mM DTT 5 mM Na<sub>4</sub>P<sub>2</sub>O<sub>7</sub> | 200 | 76 |
| 21a-b | 258-391 | 10 mM PB pH 7.4 10 mM DTT 200 mM MgCl<sub>2</sub> | 200 | 48 |
| 22a | 266-391 | 10 mM PB pH 7.4 10 mM DTT 200 mM MgCl<sub>2</sub> | 200 | 48 |
| 23a-c | 266-391 | PBS pH 7.4 10 mM DTT | 200 | 48 |
| 24a | 287-391 | 10 mM PB pH 7.4 10 mM DTT 200 mM MgCl<sub>2</sub> | 200 | 48 |
| 25a | 300-391 | 10 mM PB pH 7.4 10 mM DTT 200 mM MgCl<sub>2</sub> | 200 | 48 |
| 26a | 303-391 | 10 mM PB pH 7.4 10 mM DTT 200 mM MgCl<sub>2</sub> | 200 | 48 |
| 27a | 305-379 | 10 mM PB pH 7.4 10 mM DTT 200 mM MgCl<sub>2</sub> | 200 | 48 |
| 28a | 297-421 | 10 mM PB pH 7.4 10 mM DTT 200 mM MgCl<sub>2</sub> | 200 | 48 |
| 29a | 297-412 | 10 mM PB pH 7.4 10 mM DTT 200 mM MgCl<sub>2</sub> | 200 | 48 |
| 30a | 297-402 | 10 mM PB pH 7.4 10 mM DTT 200 mM MgCl<sub>2</sub> | 200 | 48 |
| 31a | 297-396 | 10 mM PB pH 7.4 10 mM DTT 200 mM MgCl<sub>2</sub> | 200 | 48 |
| 32a-b | 297-394 | 10 mM PB pH 7.4 10 mM DTT 200 mM MgCl<sub>2</sub> | 200 | 48 |
| 33a | 297-384 | 10 mM PB pH 7.4 10 mM DTT 200 mM MgCl<sub>2</sub> | 200 | 48 |
| 34a-b | 297-394 | 10 mM PB pH 7.4 10 mM DTT | 700 | 48 |
| 35a-d | 297-394 | PBS pH 7.4 10 mM DTT | 700 | 48 |
| 36a-c | 300 – 391 | PBS pH 7.4 10 mM DTT | 700 | 48 |
| 37a | 303-391 | PBS pH 7.4 10 mM DTT | 700 | 48 |
| 38a | 258-391 | 10 mM PB pH 7.4 10 mM DTT | 700 | 48 |
| 39a-b | 258-391 | 10 mM PB pH 7.4 10 mM DTT 5 mM Phosphoglycerate | 700 | 48 |
| 40a | 258-391 | 10 mM PB pH 7.4 10 mM DTT 300 ug/ul Heparan Sulfate | 700 | 48 |
| 41a | 258-391 | 10 mM PB pH 7.4 10 mM DTT 0.1 % NaN<sub>3</sub> | 700 | 48 |
| 42a-b | 297-408 4-pmm* | 10 mM PB pH 7.4 10 mM DTT 200 mM MgCl<sub>2</sub> | 200 | 48 |
| 43a | 297-441 4-pmm | 10 mM PB pH 7.4 10 mM DTT 200 mM MgCl<sub>2</sub> | 200 | 48 |
| 44a | 266-391 S356D | 10 mM PB pH 7.4 10 mM DTT 200 mM KCl | 200 | 48 |
| 45a | 266-391 S356D | 10 mM PB pH 7.4 10 mM DTT 200 mM NaCl | 200 | 48 |
| 46a | 0N4R | PBS pH 7.4 5 mM TCEP 50 ug/mL polyA RNA | 200 | 96 |
| 47a | 0N4R | PBS pH 7.4 5 mM TCEP 5 mM L-phosphoserine | 200 | 96 |

**DTT:** 1,4-dithiothreitol **PB:** Na<sub>2</sub>HPO<sub>4</sub>, NaH<sub>2</sub>PO<sub>4</sub>; **Dextran sulfate:** molecular weight 7-20 kDa (9011-18-1, Sigma-Aldrich); **Heparan sulfate:** 50-200 disaccharide units (57459-72-0, Sigma-Aldrich); **Poly-A RNA** (26763-19-9, Sigma-Aldrich); **PBS:** Phospho-buffered saline; **TCEP:** Tris(2-carboxyethyl) phosphine; **\*4-pmm:** four-phospho mimetic mutations: S396D S400D T403D S404D **\*PB:** Na<sub>2</sub>HPO<sub>4</sub>, NaH<sub>2</sub>PO<sub>4</sub>; **\*266/297-391:** 50:50 ratio of 266LKHQ269 (3R) and 297IKHV300 (4R) -391.

**Figure 1:**
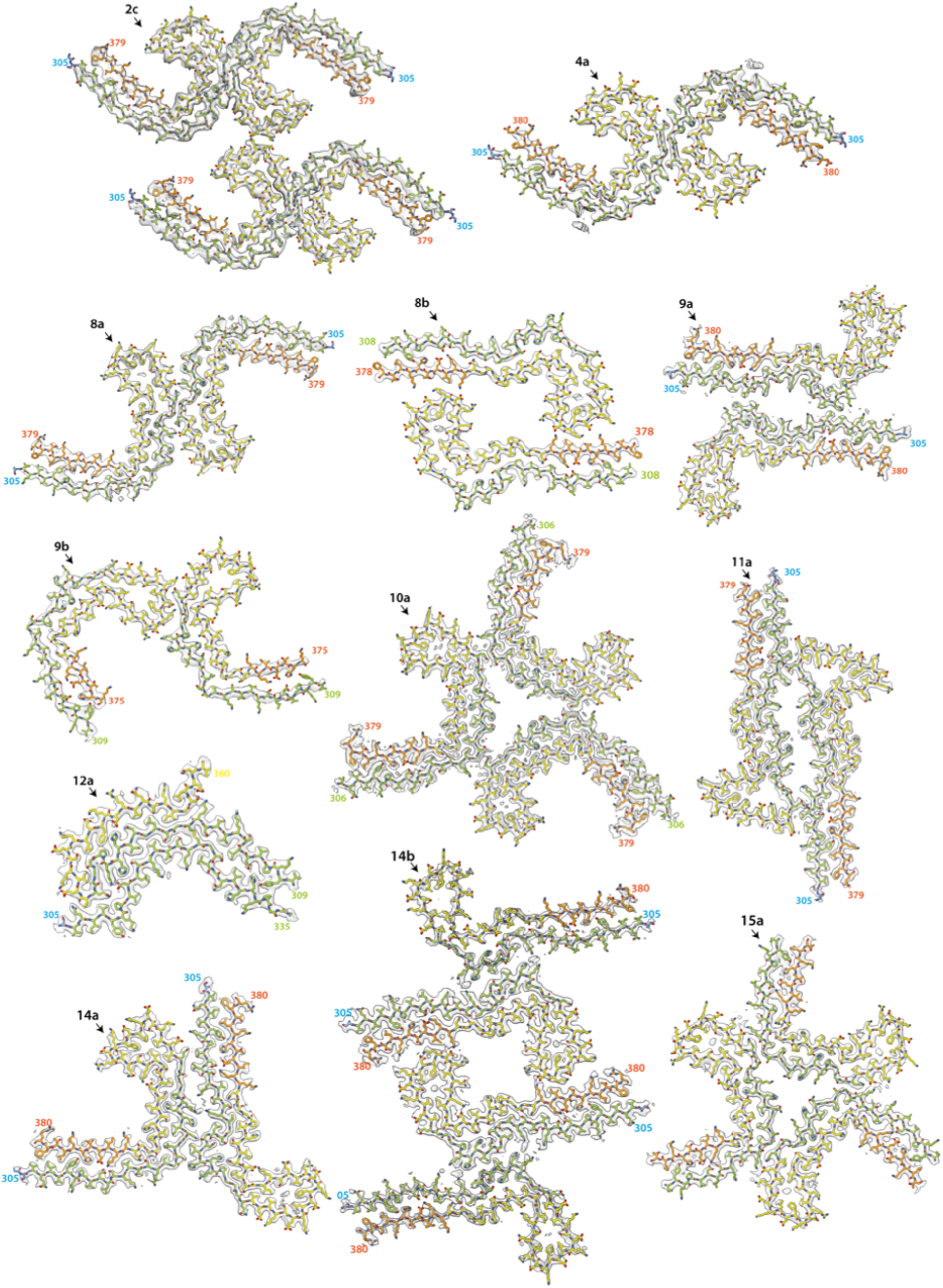
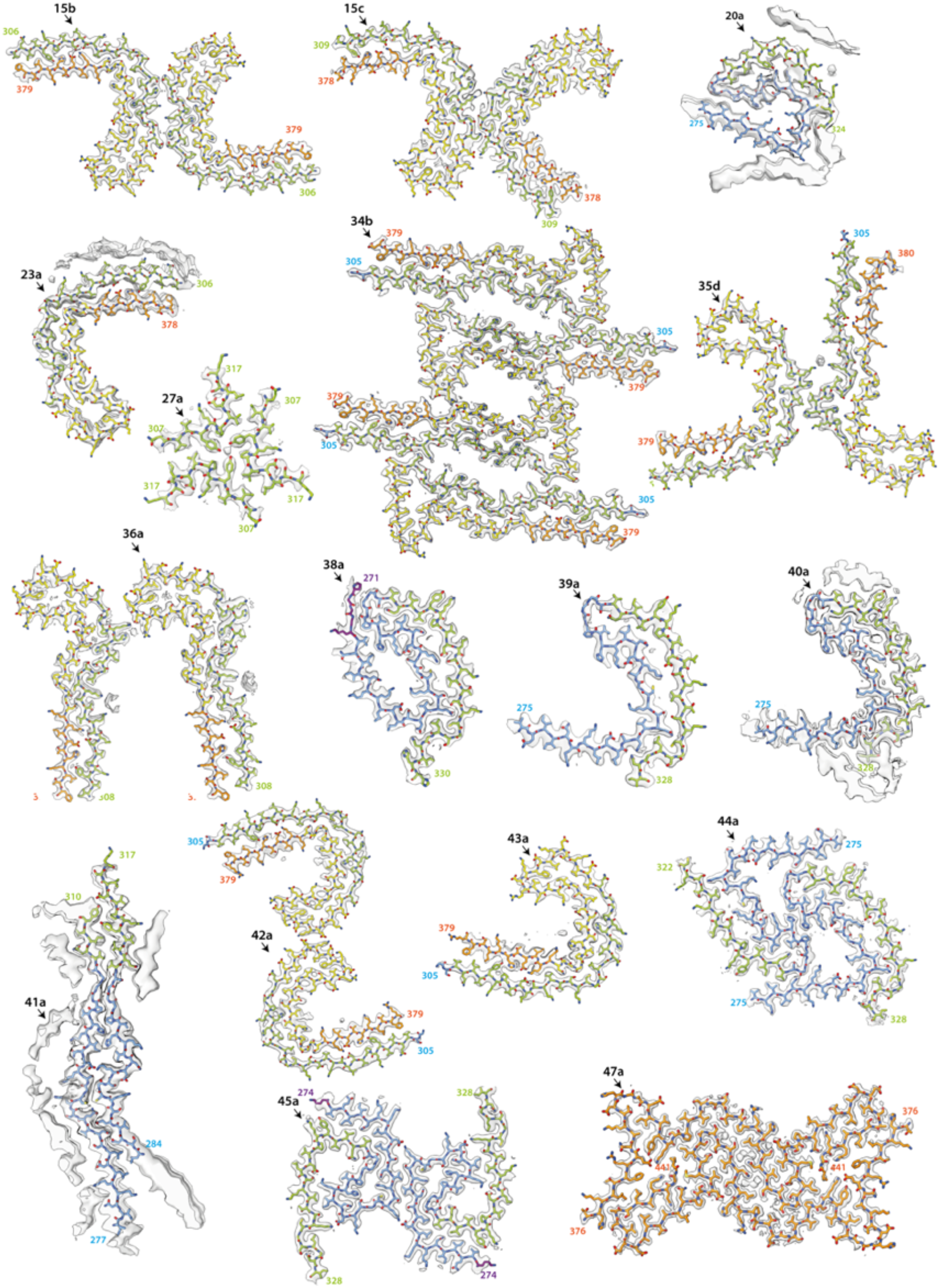
New cryo-EM structures. Cryo-EM density maps (grey transparent) and atomic models are shown for filaments with previously unobserved structures. Residues 244-274 (R1) are shown in purple; residues 275-305 (R2) are shown in blue; residues 306-336 (R3) are shown in green; residues 337-368 (R4) are shown in yellow; residues 369-441 (C-terminal domain) are shown in orange. The filament types (as defined in Table 1) are shown at the top left of each structure.

## Results

### In vitro assembly of tau(297-391) into PHFs

We first performed *in vitro* assembly of tau(297-391) in 10 mM phosphate buffer (PB) containing 10 mM dithiothreitol (DTT), with shaking at 700 rpm, as described (Al-Hilaly *et al*., 2017). Filaments formed within 4 h, as indicated by ThT fluorescence. Cryo-EM imaging after 48 h revealed a single type of filament comprising two protofilaments that were related by pseudo-2_1_ helical screw symmetry. Although the extent and topology of the ordered cores resembled those of the protofilaments of AD and CTE, the protofilament cores were extended, rather than C-shaped (**Figure 2a-c**; filament type 1a).

**Figure 2:**
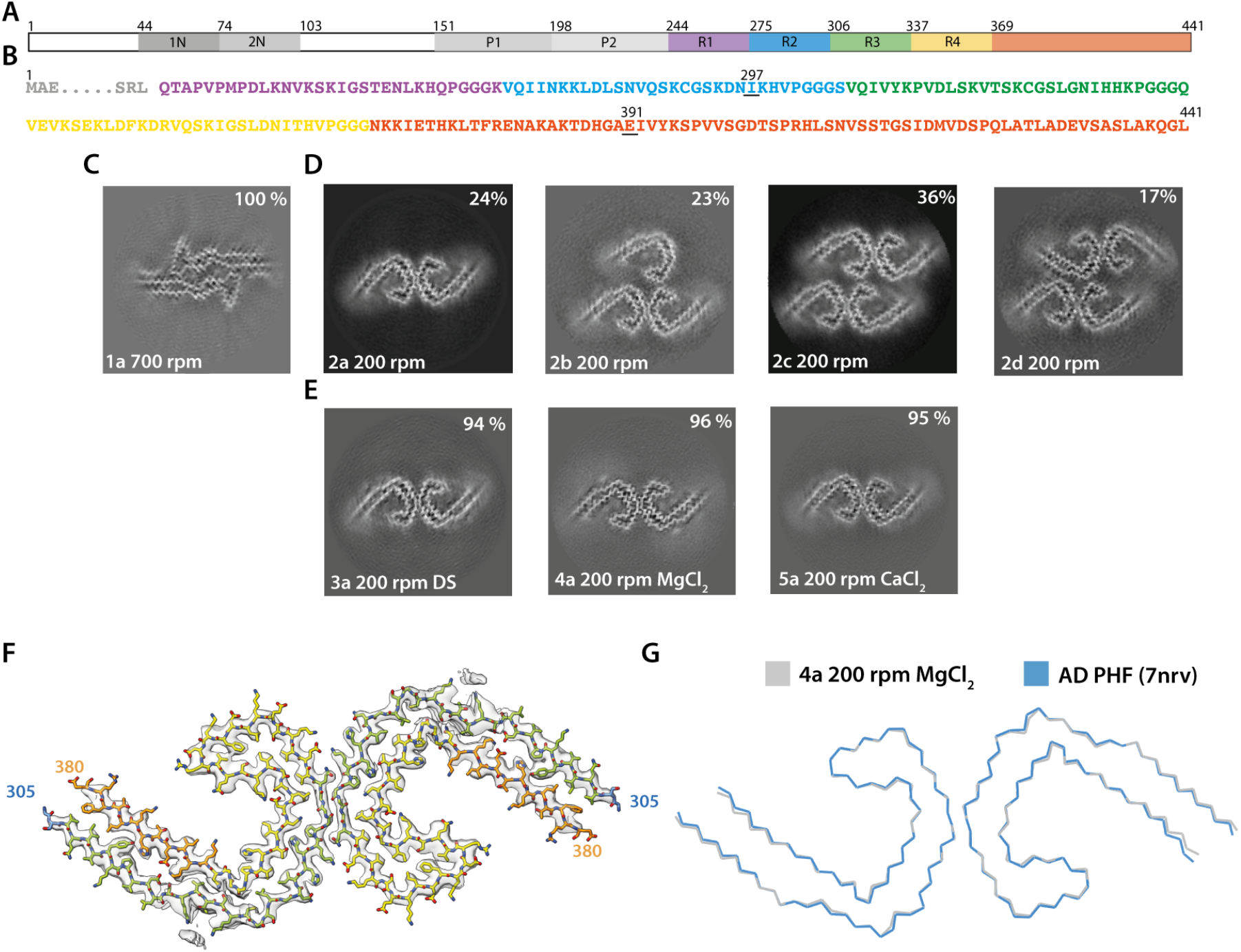
Assembly of recombinant tau into filaments like AD PHFs. **A**. Schematic of 2N4R tau sequence with domains highlighted. The regions 1N (44-73), 2N (74-102), P1 (151-197) and P2 (198-243) are shown in increasingly lighter greys; R1 (244-274) is shown in purple; R2 (275-305) is shown in blue; R3 (306-336) is shown in green; R4 (337-368) is shown in yellow; the C-terminal domain (369-441) is shown in orange. **B**. Amino acid sequence of residues 244-441 of tau, with the same colour scheme as in A. **C-E:** Projected slices, with a thickness of approximately 4.7Å, orthogonal to the helical axis for several cryo-EM reconstructions. The filament types (as defined in Table 1) are shown at the bottom left and the percentages of types for each cryo-EM data set are given at the top right of the images. **C**. Conditions of Al-Hilaly et al. (2017), with shaking at 700 rpm. **D**. Our adapted protocol, using 200 rpm shaking. From left to right; Paired Helical Filament (PHF), Triple Helical Filament (THF), Quadruple Helical Filament Type 1 (QHF-T1) and Quadruple Helical Filament Type 2 (QHF-T2) **E**. Our optimised conditions for *in vitro* assembly of relatively pure PHFs. **F**. Cryo-EM density map (grey transparent) of *in vitro* assembled tau filaments of type 4a and the atomic model colour coded according as in A. **G**. Backbone ribbon of *in vitro* PHF (grey) overlaid with AD PHF (blue).

We then reduced the shaking speed to 200 rpm. This resulted in a slower assembly reaction, with filaments appearing after 6 h. Cryo-EM structure determination after 48 h showed that the filaments were polymorphic. They shared the AD protofilament fold, spanning residues 305-378. However, besides AD PHFs, we also observed filaments comprising three or four protofilaments, which we called triple and quadruple helical filaments (THFs and QHFs) (**Figure 2d**; filament types 2b-d). THFs and QHFs have not been observed in brain extracts from individuals with AD (Fitzpatrick *et al*., 2017; Falcon, Zhang, Schweighauser, *et al*., 2018), possibly because the presence of the fuzzy coat would hinder their formation. SFs were not seen. THFs consist of a PHF and an additional single protofilament with the AD fold, whereas QHFs are made of two stacked PHFs that come in two different arrangements (types 1 and 2). The cryo-EM maps of QHF type 1 filaments were of sufficient quality to build atomic models. QHF type 1 and the THF share a common interface, where one protofilament forms a salt bridge at E342 with K343 from the third, adjoining protofilament. This protofilament remains on its own in THFs, whereas it forms a typical PHF interface with a fourth protofilament in QHFs. Although the cryo-EM reconstruction of the QHF type 2 filament was of insufficient resolution for atomic modelling, the cross-section perpendicular to the helical axis suggested that a salt bridge was present between E342 from one PHF and K321 from another (**Figure 1** and **Figure 1 - figure supplement 1;** filament types 2b-d).

Next, we sought to optimise the assembly conditions. Since THFs and QHFs have not been observed in AD, their formation would confound the use of this assembly assay for screens or to model diseases. In addition, filaments from the above experiments tended to stick together, which complicated their cryo-EM imaging, and could interfere with their use. Over longer incubation periods (> 76 h), filaments tended to precipitate, resulting in cloudy solutions. To assemble tau(297-391) into pure PHFs and reduce stickiness, we explored the addition of salts and crowding reagents. In particular, the addition of 200 mM MgCl_2_, 20 mM CaCl_2_ or 0.1 μg/ml dextran sulfate resulted in purer populations of PHFs (∼95%) (**Figure 2e**; filament types 3-5). Cryo-EM structure determination of filaments made using these conditions confirmed that their ordered cores were identical to those of AD PHFs (**Figure 2f-g**).

Tau(297-391) comprises the C-terminal 9 residues of R2, the whole of R3 and R4, as well as 23 amino acids after R4. The equivalent 3R tau construct, which lacks R2, begins with the C-terminal 9 residues of R1, the first 4 of which (^266^LKHQ^269^) are different from R2. We also assembled tau(266-391), excluding R2, in the presence of 200 mM MgCl_2_, as well as a 50:50 mixture of the 3R/4R tau constructs, in the absence of MgCl_2_. We observed PHFs, THFs and QHFs in the absence of MgCl_2_, whereas assembly in the presence of 200 mM MgCl_2_ gave rise to AD PHFs with a purity greater than 94 % (**Figure 1 - figure supplement 1;** filament types 6 and 7). These findings show that *bona fide* PHFs can be formed from only 3R or 4R, albeit truncated, tau. In AD and CTE, all six isoforms, each full-length, are present in tau filaments (Goedert *et al*., 1992; Schmidt *et al*., 2001). It remains to be determined if there are PHFs in human diseases that are made of only 3R or 4R tau.

The cryo-EM structures of *in vitro* assembled PHFs and of AD PHFs shared the same left-handed twist and pseudo-2_1_ helical screw symmetry. Moreover, there were similar additional densities in front of lysine residues 317 and 321, and on the inside of the protofilament’s C, as we previously observed for AD PHFs (**Figure 2 - figure supplement 1**). The assembly buffers contained only Na_2_HPO_4_, NaH_2_PO_4_, MgCl_2_ and DTT. Although we cannot exclude the possibility that negatively charged co-factors may have purified together with recombinant tau, it appears more likely that the additional densities arose from phosphate ions in the buffer. The phosphates’ negative charges may have counteracted the positive charges of stacked lysines. It remains to be established if similar densities in AD PHFs also correspond to phosphate ions, or if other negatively charged co-factors or parts of the fuzzy coat may play a role. The fuzzy coat, which consists of only a few residues on either side of the ordered core, is not visible in cryo-EM micrographs of *in vitro* assembled tau(297-391) (**Figure 2 – figure supplement 1)**.

### The effects of salts on tau filament assembly

During optimisation of assembly, we noticed that different cations in the buffer caused the formation of filaments with distinct protofilament folds. Besides MgCl_2_ and CaCl_2_, which led to the formation of AD PHFs, we also explored the effects of ZnCl_2_, CuCl_2_, NaCl, LiCl and KCl (**Figure 3a**). Addition of ZnCl_2_ resulted in a fold identical to that of filaments assembled using the conditions of Al-Hilaly et al. (2017), whereas addition of CuCl_2_ led to folds with little resemblance to previously observed tau folds. (**Figure 1** and **Figure 1 - figure supplement 1**; filament types 11 and 12). Cu^2+^ ions led to the formation of intermolecular disulfide bonds that were part of the ordered cores of these filaments.

**Figure 3:**
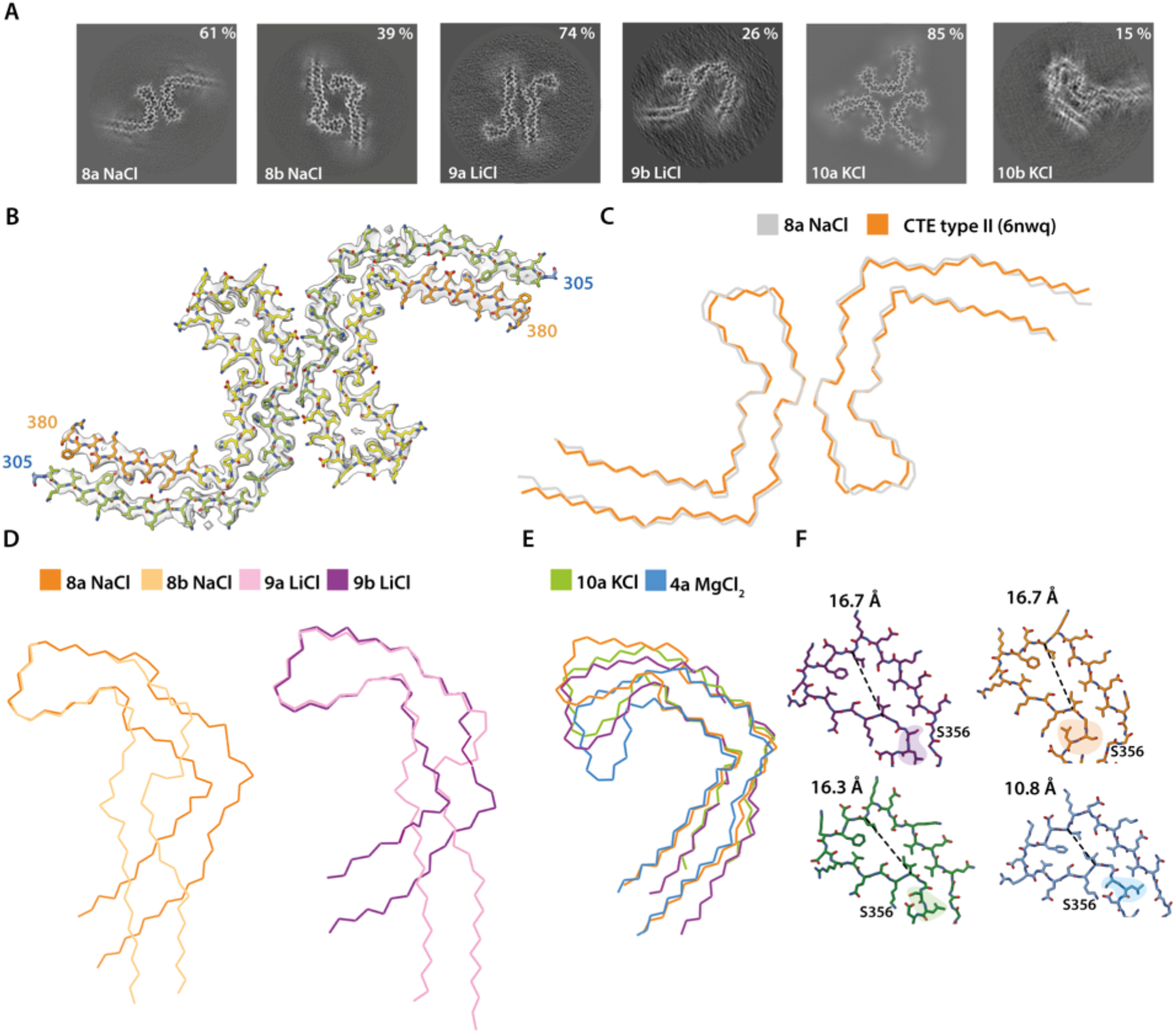
Assembly of recombinant tau into filaments like CTE type II filaments. **A**. Projected slices, with a thickness of approximately 4.7Å, orthogonal to the helical axis are shown for different assembly conditions and filament types (as defined in Table 1), which are indicated in the bottom left. The percentages of types are shown in the top right of each panel. **B**. Cryo-EM density map (grey transparent) of filament type 8a and the corresponding atomic model with the same colour scheme as in Figure 1. **C-E**. Backbone ribbon views of protofilament and filament folds. **C**. *In vitro* NaCl filament type 8a (grey) overlaid with CTE type II (orange). **D**. Extended and C-shaped protofilaments aligned at residues 338-354 for LiCl, filament types 9a and 9b (left) and NaCl, filament types 8a and 8b (right). **E**. Filament types 8b (NaCl), 9a (LiCl), 10a (KCl) and 4a (MgCl2) aligned at residues 356-364. **F**. Atomic view of residues 334-358. The distance between the Cα of L344 and I354 are indicated. Filament types 8a, 8b, 9a, 9b and 10a are shown in light purple, dark purple, dark orange, light orange and blue, respectively.

Monovalent cations modulated the formation of protofilament folds that were similar or identical to AD and CTE folds. The CTE fold is similar to the AD fold, in that it also comprises a double-layered arrangement of residues 274/305-379; however, it adopts a more open C-shaped conformation and comprises a larger cavity at the tip of the C (residues 338-354), which is filled with an additional, unknown density (Falcon *et al*., 2019). We first describe how different monovalent cations led to the formation of both C-shaped and more extended protofilament folds. We then present the effects of cations on the additional density in the cavity and the conformations of the surrounding residues.

Addition of 200 mM NaCl led to the formation of two types of filaments. The first type was identical to CTE type II filaments (**Figure 3a-c**; filament type 8a); in the second type (filament type 8b), two identical protofilaments with a previously unobserved, extended protofilament fold packed against each other with pseudo-2_1_ helical symmetry. This fold resembled the extended fold observed when using the assembly conditions of Al-Hilaly *et al*. (2017). The extended fold concurred with a flipping of the side chains of residues 322-330, which were alternatively buried in the core or solvent-exposed in opposite manner to the CTE fold. Side chains before and after ^364^PGGG^367^ had the same orientations, but formed a 90° turn in the CTE fold and adopted a straight conformation in the extended fold. Residues 338-354 had identical conformations at the tips of the C-shaped and extended folds (**Figure 1**; **Figure 3 - figure supplement 1)**. When adding 200 mM LiCl, we observed two types of filaments, with either C-shaped or more extended protofilament folds (**Figure 3a**; filament types 10a and 10b). In the first type, two C-shaped protofilaments packed against each other in an asymmetrical manner. In the second type, two protofilaments with an extended conformation packed against each other with pseudo-2_1_ helical symmetry. As observed for the filaments obtained with NaCl, the side chain orientations of residues 322-330 differed between folds. However, whereas the side chain of H330 was buried in the core of the C-shaped protofilament formed with NaCl, it was solvent-exposed in the C-shaped protofilament formed with LiCl. This suggests that the conformation of the ^364^PGGG^367^ motif defines the extended or C-shaped conformation (**Figure 3 - figure supplement 2**). Addition of 200 mM KCl also led to two different filaments with either extended or C-shaped protofilament folds. However, in this case, low numbers of filaments with extended protofilaments resulted in poor cryo-EM reconstructions. The filaments with C-shaped folds comprised three protofilaments, which packed against each other with C3 symmetry (**Figure 1; Figure 3a**; filament type 11a).

For each monovalent cation, residues 338-354 adopted identical conformations when comparing extended and C-shaped protofilament folds (**Figure 3d**). These residues surrounded the cavity at the tip of the fold, which was filled with an additional density in the CTE fold. Additional densities were also observed in filaments formed in the presence of NaCl, KCl, and in the extended filaments formed with LiCl. The cryo-EM reconstructions of the 3-fold symmetric filaments formed with KCl, with a resolution of 1.9 Å, showed multiple additional spherical densities inside the protofilament core. Besides additional densities for what were probably water molecules in front of several asparagines and glutamines, the cavity at the tip of the fold contained two larger, separate spherical densities per β-rung, which were 3.1 Å apart, and at approximately 3-4.5 Å distance from S341 and S352, the only polar residues in the cavity. Another pair of additional densities, similar in size to those inside the cavity, was present at a distance of 2.6 Å from the carbonyl of G335 (**Figure 3 – figure supplement 3**). Below, we will argue that these densities corresponded to pairs of K^+^ and Cl^-^ ions. Reconstructions for the filaments formed with NaCl were at resolutions of 2.8 and 3.3 Å. The additional density in these maps was not separated into two spheres, but was present as one larger blob per rung, with separation between blobs along the helical axis. Filaments formed with LiCl were resolved to resolutions of 3.1 and 3.4 Å. No additional densities were present inside the cavity of the C-shaped fold, but the cavity in the extended fold contained a spherical density that was smaller than the densities observed for NaCl and KCl filaments (**Figure 3 – figure supplement 3**). Different cations also led to conformational differences in residues S356 and L357, which were akin to the differences observed previously between AD and CTE folds (**Figure 3e-f)** (Falcon *et al*., 2019). In the AD fold, S356 is solvent-exposed and L357 is buried inside the protofilament core, whereas they adopt opposite orientations in the CTE fold. As mentioned, filaments formed with NaCl are identical to CTE filaments; in filaments formed with KCl, S356 and L357 are oriented in the same directions as in the AD fold; in filaments formed with LiCl, both residues are buried in the core.

Efforts to generate AD SFs or CTE type I filaments using mixtures of salts were unsuccessful, but they did reveal novel protofilament interactions (**Figure 1; Figure 1 - figure supplement 1**; filament types 13-17). Notably, using a buffer with MgSO_4_ and NaCl, we obtained a minority of filaments with an SF interphase (11 %, filament type 15d). Probably because of the presence of NaCl, protofilaments adopted the CTE fold. Further exploration of the role of salts may lead to the assembly of recombinant tau into AD SFs and CTE type I filaments.

### The effects of protein length on tau filament assembly

We investigated if the N- and C-termini of tau(297-391) are required for its assembly into PHFs. We first made a series of protein fragments ending at residue 391, to explore the effects of the position of the N-terminus. Next, keeping the N-terminus at residue 297, we explored the effects of the position of the C-terminus (**Figure 4a**). Each recombinant tau fragment was assembled in 10 mM PB, 10mM DTT and 200 mM MgCl_2_, at 200rpm shaking. We assessed the presence of tau filaments by negative stain EM and used cryo-EM to determine their structures (**Figure 4b**; filament types 18-30).

**Figure 4:**
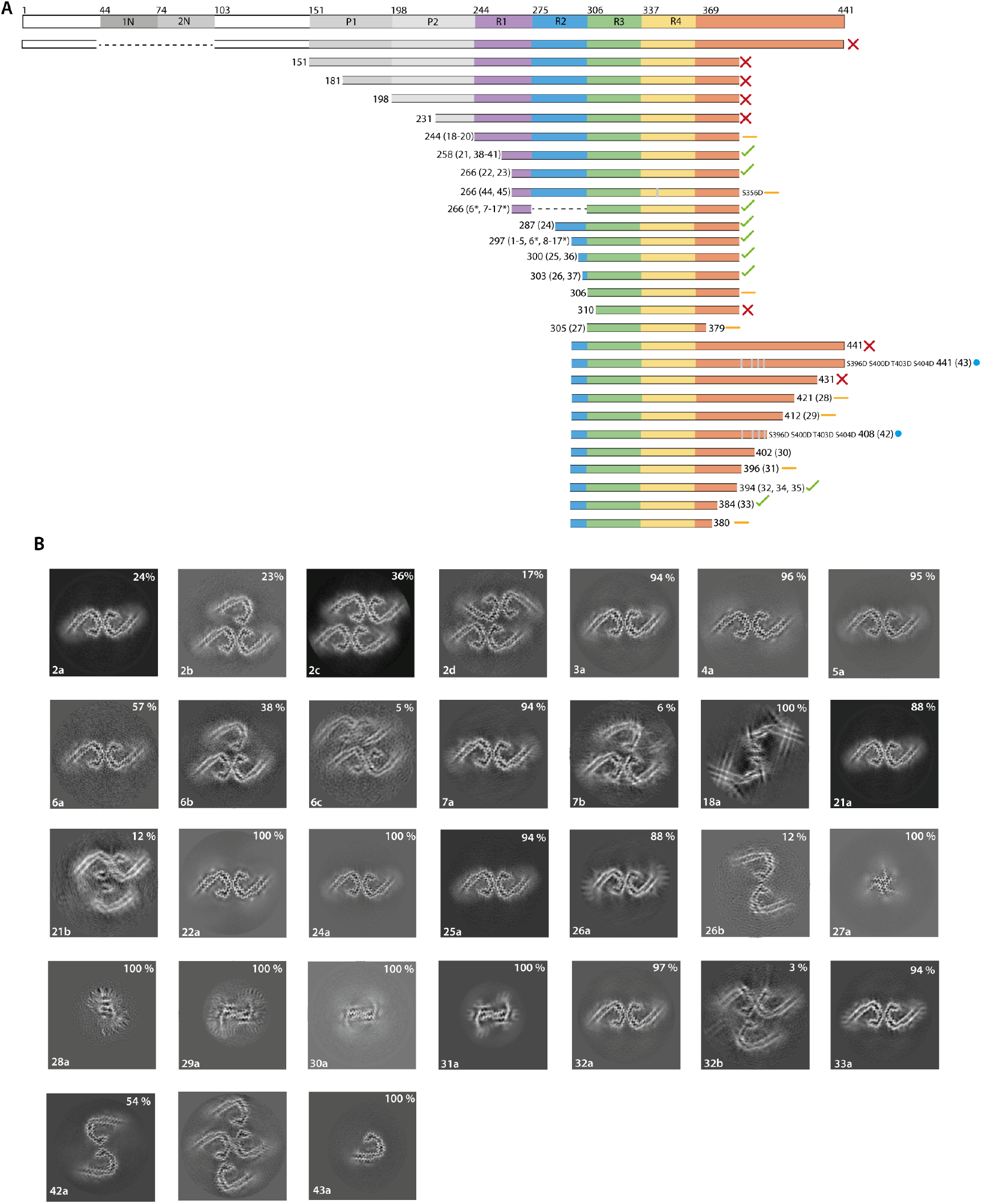
The effects of protein length on tau filament assembly. **A**. Schematic representation of 2N4R tau and the constructs used in this study. A red cross indicates that no filaments were formed; an orange dash indicates that filaments with structures distinct from AD PHFs were formed; a blue circle indicates that the AD protofilament fold was formed; a green tick indicates that AD PHFs were formed. Resulting filament types (as defined in Table 1) are indicated for each experiment. **B**. Projected slices, with a thickness of approximately 4.7Å, orthogonal to the helical axis for the filaments formed with the constructs in A. Filament types are indicated at the bottom left; percentages of filament types in each cryo-EM data set are shown in the top right. (*) 50:50 ratio of 266-391 (3R) and 297-391 (4R) tau.

Proteins comprising the entire N-terminal domain (residues 1-391 of 0N4R tau) did not assemble into filaments. The same was true of proteins starting at residues 151, 181 or 231 in the proline-rich region. When using proteins starting at 244, the first residue of R1, we observed the formation of filaments. The quality of the cryo-EM reconstructions was not sufficient for atomic modelling, but the ordered filament cores adopted a more open conformation than in the Alzheimer fold, with two protofilaments interacting at their tips. Residues 338-354 probably adopted the same conformation as in the Alzheimer fold. Addition of NaCl or pyrophosphate did not lead to the formation of PHFs or CTE filaments (**Figure 1; Figure 1 – figure supplement 1**; filament types 19-20). However, it is possible that this construct will form filaments with the Alzheimer fold after further optimisation of assembly conditions. Proteins that started at residues 258 or 266 in R1 formed PHFs. Adding NaCl to tau(297-391), or using phosphate-buffered saline (PBS), gave rise to CTE type 2 filaments, as well as to filaments consisting of either single CTE protofilaments or two protofilaments, related by C2 symmetry, which formed an interface at residues 327-336 (**Figure 1 - figure supplement 1**; filament types 15b, 23b). For all filaments assembled in the presence of NaCl, we saw densities inside the tip of the fold’s cavity. Moreover, with the proteins starting at residues 258 and 266, we observed proteinaceous densities, which packed against the N-terminus of the protofilaments. This may have been the extension into R2 (**Figure 1 - figure supplement 1**; filament types 23a-c).

Tau(297-391) can also be shortened from the N-terminus, as proteins starting at residues 300 or 303 still formed filaments with AD or CTE folds (**Figure 1 – figure supplement 1**; filament types 25-26). When shortened to residue 306, we also observed filaments, but their stickiness precluded cryo-EM structure determination. No filaments were formed when shortening the protein to residue 310. It has previously been shown that the PHF6 motif (^306^VQIVYK^311^ in tau) is essential for filament formation (von Bergen *et al*., 2000; Li and Lee, 2006).

Reminiscent of what we observed for the N-terminal domain, the tau starting at residue 297 and ending at 441 failed to assemble into filaments. The same was true of tau(297-431). However, proteins ending at residues 421, 412, 402 or 396 formed filaments, but their ordered cores were much smaller than in the Alzheimer fold, precluding cryo-EM structure determination to sufficient resolution for unambiguous atomic modelling (**Figure 4b;** filament types 28-31). We did observe filaments with the Alzheimer fold for a protein ending at residue 394 (**Figure 1 – figure supplement 1; Figure 4b**; filament type 32).

As observed for proteins ending at residue 384, when tau(297-391) was shortened from the C-terminus, it could still form AD and CTE folds (**Figure 4b**; filament type 33). However, constructs ending at residue 380 formed straight ribbons, precluding cryo-EM structure determination.

As shown in Figure 2, we found that reducing the shaking speed from 700 to 200 rpm was important for the formation of filaments with the AD fold by tau(297-391). To show that this was not only the case for tau(297-391), we assembled tau(258-391) and tau(297-394) with a shaking speed of 700 rpm. For tau(297-394), besides structures similar to those formed by tau(297-391) at 700 rpm, we observed an additional filament with pseudo-2_1_ helical screw symmetry that comprised 4 protofilaments (**Figure 1; Figure 1 - figure supplement 1**; filament type 34). In the presence of NaCl, we observed filaments with similar extended conformations, but with a CTE cavity. This was also the case when tau(300-391) and tau(303-391) were assembled at 700 rpm in PBS **(Figure 1; Figure 1 – figure supplement 1**; filament types 35-37). When assembling tau(258-391) at 700 rpm, we observed protofilaments with partial similarity to the GGT fold. The structures were nearly identical at residues 288-322 (**Figure 1; Figure 1 – figure supplement 1; Figure 4 - figure supplement 1;** filament type 38a). It is possible that the absence of a non-proteinaceous co-factor, which was hypothesised to be present inside the GGT fold, precluded formation of *bona fide* GGT filaments. Addition of heparan sulfate or phosphoglycerate to the assembly buffer did not result in formation of the GGT fold **(Figure 1; Figure 4 – figure supplement 1**; filament types 39-41). Further experiments are needed to identify the co-factor of GGT.

### The effects of pseudophosphorylation on tau filament assembly

Whereas *in vitro* filament assembly of recombinant tau repeats was inhibited by N- and C-terminal regions, it is full-length tau that assembles into filaments in AD (Lee *et al*., 1991; Goedert *et al*., 1992). This difference may be due to the fact that we used unmodified recombinant proteins for *in vitro* assembly, while tau undergoes extensive post-translational modifications in AD (Wesseling *et al*., 2020). Immunochemistry and mass spectrometry have identified abnormal hyperphosphorylation of S396, S400, T403 and S404 in PHF-tau (Lee *et al*., 1991; Hasegawa *et al*., 1992; Kanemaru *et al*., 1992; Morishima-Kawashima *et al*., 1995). Because these sites are located in the C-terminal domain that prevents the formation of PHFs from recombinant tau, we hypothesised that their phosphorylation may modulate PHF formation and help to overcome the inhibitory effects of the C-terminal domain. We therefore mutated these residues to aspartate, with its negative charge mimicking the negative charges of phosphorylation, in the tau constructs ending at residues 408 and 441 (**Figure 4**; filament types 42-43).

As described above, tau(297-408) formed filaments with small ordered cores; its pseudophosphorylated mutant formed two types of filaments consisting of AD PHF protofilaments. Both filament types had the same inter-protofilament interface as that of THFs and type 1 QHFs, where E342 from one protofilament forms a salt bridge with K343 of the other. The second filament type was a doublet of the first. Importantly, tau(297-441), which does not assemble into filaments, did so in presence of the four phospho-mimetic mutations. Filaments consisted of a single protofilament with the AD fold.

It has been suggested that AD filaments may also be phosphorylated at S356 (Hanger *et al*., 1998, 2007). We therefore tested phospho-mimetic mutation S356D. However, using tau(266-391) with S356D, we were unable to form PHFs. Instead, filaments consisted of two asymmetric protofilaments, both of which had features reminiscent of the GGT fold. In the presence of NaCl, this construct formed filaments with a novel fold, consisting of two identical protofilaments related by pseudo-2_1_ helical symmetry (**Figure 1, Figure 1 - figure supplement 1** and **Figure 4 - figure supplement 1**; filament types 44 and 45).

### The effects of anionic co-factors on tau filament assembly

Addition of anionic co-factors leads to the formation of filaments from full-length tau (Goedert *et al*., 1996; Kampers *et al*., 1996; Pérez *et al*., 1996; Wilson and Binder, 1997; Farid, Corbo and Alonso, 2014). However, we previously showed that heparin-induced tau filament formation leads to structures that are different from those present in disease (Zhang *et al*., 2019). To further explore the effects of anionic co-factors, we solved cryo-EM structures of filaments formed using full-length recombinant tau in the presence of RNA (Kampers *et al*., 1996) or phosphoserine. The resulting tau filaments had structures that have not been observed previously.

Addition of RNA led to the formation of two asymmetrical and extended protofilaments, similar to heparin-induced 3R tau filaments (Zhang *et al*., 2019) (**Figure 1 - figure supplement 1;** filament type 46). The resulting map was of insufficient resolution for atomic modelling. We calculated a 1.8 Å resolution reconstruction for full-length tau filaments that formed in the presence of 5 mM phosphoserine **(Figure 1 - figure supplement 1**; filament type 47). At this resolution, the absolute handedness of the filament was obvious from the position of the main-chain carbonyl oxygens. Whereas all previously described tau filaments had a left-handed twist, phosphoserine-induced filaments were right-handed. Surprisingly, the ordered core of this filament comprised residues 375-441, encompassing only residues from the C-terminal domain. Filaments consisted of two protofilaments that were related by pseudo-2_1_ helical symmetry and the protofilament fold contained 8 β-sheets. Several water molecules were observed, particularly in front of the side chains of serines, threonines and asparagines. This is the first example of a region outside the microtubule-binding repeats of tau forming amyloid filaments.

## Discussion

In the course of protein evolution, natural selection has produced amino acid sequences that fold into specific protein structures to fulfil the multitude of tasks required to maintain life. The observation that the same proteins can adopt multiple different amyloid structures has made it clear that the paradigm by which a protein’s sequence defines its structure may not hold for amyloids. Whereas the folding of globular proteins is mainly driven by hydrophobic collapse, amyloid filament formation is driven by hydrogen bond formation between the extended β-sheets along the helical axis (Fitzpatrick *et al*., 2011). Apparently, the packing of β-sheets against each other can happen in many ways for a single amino acid sequence, since multiple different amyloid structures have been reported for proteins like tau (Zhang *et al*., 2019; Shi *et al*., 2021), amyloid-β (Kollmer *et al*., 2019; Yang *et al*., 2021), α-synuclein (B. Li *et al*., 2018; Y. Li *et al*., 2018; Guerrero-Ferreira *et al*., 2019; Schweighauser *et al*., 2020; Lövestam *et al*., 2021), TAR DNA binding protein-43 (TDP-43) (Arseni *et al*., 2021; Li, Babinchak and Surewicz, 2021) and immunoglobulin light chain (Radamaker *et al*., 2019; Swuec *et al*., 2019). Because amyloid filaments of these proteins are typically associated with pathology, rather than function, their structural diversity could be disregarded. However, the observation that, for tau, and possibly also other proteins, the different amyloid structures define distinct neurodegenerative conditions, raises important questions on what drives their formation.

The lack of *in vitro* assembly models for recombinant tau that replicate the amyloid structures observed in disease has hampered progress. Here, we identified conditions for the *in vitro* assembly of AD PHFs and type II CTE filaments, and established the formation of these structures by cryo-EM structure determination to resolutions sufficient for atomic modelling. The latter is crucial. Biochemical methods that discriminate between the different structures in solution do not exist and negative stain EM or atomic force microscopy do not provide sufficient resolution to unequivocally distinguish between them. Even distance-restraints derived from solid-state NMR are probably not sufficient to tell all the different structures apart, in particular in terms of inter-molecular interactions, such as protofilament packing.

The ability to make AD PHFs and CTE type II filaments *in vitro* opens up new avenues for studying tauopathies. *In vitro* assembly assays could be used to screen for compounds that inhibit filament formation. Alternatively, *in vitro* generated filaments may be used in high-throughput screens for small-molecule compounds or biologics that bind specifically to a single type of filament. Such amyloid structure-specific binders could be developed into ligands for positron emission tomography to differentiate between tauopathies in living patients. Moreover, specific binders could be explored for use in therapeutic approaches that aim to degrade filaments inside neurons through their coupling with the protein degradation machinery inside the cell. Ultimately, specific binders could even obviate the need for cryo-EM structure determination to confirm the formation of the correct types of filaments in new model systems for disease.

Besides their use in screens, *in vitro* assembly of tau filaments also provides a model system, amenable to experimental perturbation, for the study of the molecular mechanisms that underlie amyloid formation. Coupled to cryo-EM structure determination, this provides a promising model system for studying the formation of different tau folds that define distinct tauopathies. The work presented in this paper provides two examples, as outlined below.

### Ideas and speculations

A major difference between Alzheimer and CTE tau folds is the presence of a larger cavity that is filled with an additional density in the CTE fold. Based on the relatively hydrophobic nature of this cavity, we previously hypothesised that this additional density may correspond to an unknown hydrophobic co-factor that assembles with tau to form CTE filaments (Falcon *et al*., 2019). However, we now observe the *in vitro* assembly of CTE type II filaments in the absence of hydrophobic co-factors. Instead, whether CTE type II filaments or AD PHFs formed *in vitro* was determined by the presence or absence of NaCl. Moreover, using different monovalent cations changed the additional densities, as well as the conformation of residues surrounding the cavity. The 1.9 Å map for the structure formed with KCl showed two spherical blobs of additional density per β-rung that were in an arrangement that would be consistent with a pair of K^+^ and Cl^-^ ions; a similar pair of spherical densities was also observed in front of G335 (**Figure 3 - figure supplement 3**). Therefore, it is possible that the cavity in the filaments formed with NaCl is filled with Na^+^ and Cl^-^ ions. The continuous nature of the additional density along the helical axis could arise from limited resolution of the reconstructions with NaCl, or from the ions not obeying the 4.75 Å helical rise that is imposed on the reconstruction. The observation that the filaments formed with NaCl are identical to CTE type II filaments suggests that, rather than being filled with hydrophobic co-factors, the cavity in CTE filaments may contain NaCl. Na^+^ and Cl^-^ levels in neurons are typically much lower than the 200 mM NaCl used here, but it could be that brain trauma somehow leads to increased levels of these ions in the brain regions where tau filaments first form. Additional studies will explore this further.

Our results have also identified how regions outside the repeats interfere with the *in vitro* assembly of AD PHFs. In particular, the presence of the N-terminal domain, including its proline-rich region, and residues beyond 421 in the C-terminal domain, inhibited the spontaneous assembly of recombinant tau. In addition, the presence of residues 396-421 led to the formation of filaments with much smaller ordered cores. The C-terminal region has been reported to inhibit anionic co-factor-induced assembly of full-length tau, with pseudophosphorylation overcoming inhibition (Abraha *et al*., 2000; Haase *et al*., 2004).

Interestingly, mutating serine or threonine residues at positions 396, 400, 403 and 404 to aspartate, to mimic phosphorylation, overcame these inhibitory effects. Although we did not yet examine the effects of pseudophosphorylation of the N-terminal domain, we note that the proline-rich region upstream of the repeats is positively charged, which may inhibit amyloid formation. The introduction of negative charges through phosphorylation, or the removal of positive charges by, for instance, acetylation, may be necessary for filament assembly. In AD, filamentous tau is extensively hyperphosphorylated, especially in the proline-rich region and the C-terminal domain (Grundke-Iqbal *et al*., 1986; Iqbal, Liu and Gong, 2016). Hyperphosphorylation inhibits the ability of tau to interact with microtubules (Bramblett *et al*., 1993; Yoshida and Ihara, 1993), which may be necessary for filament formation, since the physiological function of tau and its pathological assembly require the repeat region. Other post-translational modifications, such as acetylation and ubiquitination of positively charged residues in the microtubule-binding repeats, may also play a role (Wesseling *et al*., 2020). It is therefore possible that post-translational modifications of tau lead to a higher propensity to form amyloid filaments in disease. Thus, the enzymes causing these modifications could be therapeutic targets.

Moreover, most of the structures in **Figure 1**, all of which are of recombinant tau proteins over-expressed in *E*.*coli*, share features with the Alzheimer and CTE folds: a cross-β packing of residues from near the start of R3 against residues from near the start of the C-terminal domain, and a turn in R4. Other tau folds are markedly different. Unmodified tau may have a tendency to form filaments that resemble Alzheimer and CTE folds, whereas specific post-translational modifications and/or associated molecules may play a role in driving the formation of tau folds associated with other tauopathies.

## Outlook

For many years, amyloid formation has been studied in laboratory-based model systems for disease. Until recently, these studies were blind to the structures being formed. Our work illustrates how cryo-EM structure determination can guide the development of better experimental models by showing that the filaments generated from recombinant tau are identical to those formed in disease. As a start, we identified *in vitro* assembly conditions for the formation of tau filaments like those of AD and CTE. Future work will explore which factors, including post-translational modifications and/or associated molecules, determine the formation of other tau filaments.

Our results do not only have implications for the *in vitro* assembly of tau. Similar strategies could be applied to the study of other amyloid-forming proteins, and insights from *in vitro* assembly studies may carry over to the development of better experimental models of disease, including in cell lines and in animals. Given the structural diversity of amyloids, we envision that high-throughput cryo-EM structure determination will play a crucial role in future developments.

## Author information

## Acknowledgements

We thank R.A. Crowther, J.G. Greener and M. Wilkinson for helpful discussions; T. Darling and J. Grimmett for help with high-performance computing; D. Cats and M. van Beers for help with the Glacios and Krios G4 systems at Eindhoven; and the Electron Microscopy Facility of the MRC Laboratory of Molecular Biology and the Thermo Fisher Scientific Research and Development Facility for help with cryo-EM data acquisition. This work was supported by the U.K. Medical Research Council (MC-U105184291 to M.G. and MC_UP_A025_1013, to S.H.W.S.).

## Author contributions

S.L. performed experiments and cryo-EM data acquisition, except for cryo-EM data acquisition in Eindhoven, which was performed by F.A.K. and A.K.; B.v.K. helped with the EPU multigrid algorithm; S.L., A.G.M. and S.H.W.S. analysed cryo-EM data; S.H.W.S. and M.G. supervised the project; all authors contributed to the writing of the manuscript.

## Data and materials availability

There are no restrictions on data and materials availability. Cryo-EM maps and atomic models have been deposited at the EMDB and the PDB, respectively (see **Supplementary Tables 1-24**).

## Methods

### Cloning

Constructs were made with *in vivo* assembly (IVA) cloning (García-Nafría, Watson and Greger, 2016) using pRK172 0N4R human tau. Reverse and forward primers were designed to share 15-20 nucleotides of homologous region and 15-30 nucleotides for annealing to the template with melting temperatures ranging from 58 - 67 °C.

### Protein Expression and Purification

Expression of tau was carried out in *E. coli* BL21 (DE3) gold cells, as described (Studier, 2005). In brief, a single colony was inoculated into 500 ml lysogeny broth (LB) auto-induction media and grown for 8 h at 37 °C, followed by subsequent expression for 16h at 24 °C. Cells were harvested by centrifugation (4,000 x g for 20 min at 4 °C), and resuspended in washing buffer (WB: 50 mM MES at pH 6.0; 10 mM EDTA; 10 mM DTT, supplemented with 0.1 mM PMSF and cOmplete EDTA-free protease cocktail inhibitors, at 10 ml/g of pellet). Cell lysis was performed using sonication (at 40% amplitude using a Sonics VCX-750 Vibracell Ultra Sonic Processor for 7 min, 5 s on/10 s off). Lysed cells were centrifuged at 20,000 x g for 35 min at 4 °C, filtered through 0.45 μm cut-off filters and loaded onto a HiTrap SP HP 5 ml column (GE Healthcare) for cation exchange. The column was washed with 10 column vol of WB and eluted using a gradient of WB containing 0 – 1 M NaCl. Fractions of 3.5 ml were collected and analysed by SDS-PAGE bis-TRIS 4-12 % or Tris Glycine 4-20%. Protein-containing fractions were pooled and precipitated using 0.3 g/ml ammonium sulphate and left on a rocker for 30 min at 4 °C. Precipitated proteins were then centrifuged at 20,000 x g for 35 min at 4 °C, and resuspended in 2 ml of 10 mM phosphate buffer (PB), pH 7.2-7.4, with 10 mM DTT, and loaded onto a 60/10 Superdex size exclusion column. Size exclusion fractions were analysed by SDS-PAGE, protein-containing fractions pooled and concentrated to 6 mg/ml using molecular weight concentrators with a cut-off filter of 3 kDa. Purified protein samples were flash frozen in 100 μl aliquots for future use.

### Filament Assembly

Purified protein samples were thawed and diluted to 4 mg/ml. Filaments were assembled in a FLUOstar Omega (BMG Labtech, Aylesbury, UK) using Corning 96 Well Black Polystyrene Microplate (Thermofischer) with orbital shaking at 37 °C. See **Table 1** for duration and speed settings. The addition of compounds proceeded by first adding proteins, diluted with 10 mM PB and 10 mM DTT; salts and/or cofactors were added last. Each condition was run in two separate wells. In one well, 3 μM ThT was added to allow continuous monitoring of protein assembly. The other well, without ThT, was used for microscopy. The presence of filaments was also assessed using negative stain electron microscopy on a Thermo Fisher Scientific Tecnai Spirit (operating at 120 kV).

### Electron Cryo-Microscopy

Samples with confirmed filaments were centrifuged at 3,000 x g for 2 min to remove any large aggregates, and 3 μl aliquots were applied to glow-discharged R1.2/1.3, 300 mesh carbon Au grids that were plunge-frozen in liquid ethane using a Thermo Fisher Scientific Vitrobot Mark IV. Cryo-EM data were acquired at the MRC Laboratory of Molecular Biology (LMB) and at the Research and Development facility of Thermo Fisher Scientific in Eindhoven (TFS). All images were recorded at a dose of 30-40 electrons per Å^2^, using EPU software (Thermo Fisher Scientific), and converted to tiff format using relion_convert_to_tiff prior to processing. See **Table 2** and **Supplementary Tables 1-24** for detailed data collection parameters.

**Table 2:** Data acquisition details for the different microscopes.

| **Microscope** | LMB Krios G1 | LMB Krios G2 | TFS Glacios | TFS Krios G4 |
| --- | --- | --- | --- | --- |
| Magnification | 105,000 | 96,000 | 165,000 | 165,000 |
| Camera | K2*/K3 | Falcon 4 | Falcon 4 | Falcon 4 |
| Energy filter (eV) | 20 | NA | 20 | NA |
| Voltage (kV) | 300 | 300 | 200 | 300 |
| Electron exposure (e-/Å<sup>2</sup>) | 40 | 30/40 | 40 | 40 |
| Defocus range (µm) | 1.5-3 | 1.2-2.5 | 0.6-1.2 | 0.6-1.2 |
| Pixel size (Å) | 0.85/1.145* | 0.824 | 0.672 | 0.727 |

\* LMB Krios G1 initially had a K2 camera, which was later replaced by a K3 camera. Only condition 1a (as defined in Table 1) was collected on the K2 camera. The pixel size on the K2 camera was 1.145 Å.

At LMB, images were recorded on a Krios G1 with a K2 or K3 camera (Gatan), using an energy slit of 20 eV on a Gatan energy filter, or on a Krios G2 with a Falcon 4 camera (Thermo Fisher Scientific), without an energy filter. At TFS, images were recorded on a Krios G4 with a CFEG, a Falcon 4 camera, and a Selectris X energy filter that was used with a slit width of 10 eV.

At TFS, cryo-EM grid screening and part of data acquisition were performed using EPU-Multigrid on a Glacios microscope equipped with a Selectris X energy filter (Thermo Fisher Scientific) and a Falcon 4 camera. Eight grids were loaded for each 48 hr EPU-Multigrid run. Each grid was loaded to the stage for session set-up, grid square selection and ice-filter preparation. The session information was stored in EPU and associated with the grid position in the autoloader. After all the grids sessions were created and stored in queue, two-fold astigmatism was corrected and the beam tilt was adjusted to the coma-free axis using automatic functions within EPU. Before starting the EPU Multigrid queue, and once for each 48hr session, the Selectris X filter slit was cantered, and the filter tuned for isochromaticity, magnification and chromatic aberrations, using Sherpa software (Thermo Fisher Scientific). During the fully automated EPU Multigrid runs, each grid was loaded onto the stage, grid squares were brought to eucentric height, and holes were selected with the stored ice filter settings. For each grid, 2000 images were collected with a 10eV energy filter slit width. Images were collected in EER mode using the aberration-free image shift method in EPU version 2.12. Under these conditions, a throughput of 200-250 images per hour was achieved depending on the number of holes available per grid square.

### Helical Reconstruction

Movie frames were gain-corrected, aligned and dose weighted using RELION’s motion correction program (Zivanov, Nakane and Scheres, 2019). Contrast transfer function (CTF) parameters were estimated using CTFFIND-4.1 (Rohou and Grigorieff, 2015). Helical reconstructions were performed in RELION-4.0 (He and Scheres, 2017; Kimanius *et al*., 2021). Filaments were picked manually or automatically using Topaz (Bepler *et al*., 2019). The neural network in Topaz was trained using a subset of 10,000 extracted segments from the dataset of construct 305-379 (see table X). This trained model was used to predict positions for filaments for all other datasets using a Topaz threshold value ranging from -4 to -7, combined with custom-made Python code to output start-end coordinates of helical segments, rather than individual coordinates for all segments. The picked particles were extracted in boxes of either 1028 or 768 pixels, and then down-scaled to 256 or 128 pixels, respectively. For all data sets, reference-free 2D classification was performed to assess different polymorphs, cross-over distances and to select suitable segments for further processing. Initial models were generated *de novo* from 2D class average images using relion_helix_inimodel2d (Scheres, 2020). Subsequently, 3D classification was used to select particles leading to the best reconstructions, and 3D auto-refinement was used to extend resolution and to optimise the helical twist and rise parameters. For all rendered maps in Figures 1, 2 and 4, Bayesian polishing (Zivanov, Nakane and Scheres, 2019) and CTF refinement (Zivanov, Nakane and Scheres, 2020) were performed to further increase resolution. Final maps were sharpened using standard post-processing procedures in RELION and reported resolutions were estimated using a threshold of 0.143 in the Fourier Shell Correlation (FSC) between two independently refined half-maps (**Figure 1 - figure supplement 2**) (Chen *et al*., 2013).

### Model Building

All atomic models were built *de novo* in COOT, using 3 rungs for each structure. Coordinate refinement was performed in ISOLDE (Croll, 2018; Casañal, Lohkamp and Emsley, 2020). Dihedral angles from the middle rung, which was set as a template, were also applied to the rungs below and above. For each refined structure, separate model refinements were performed on the first half map, after increasing the temperature to 300 K for 1 min. The resulting model was then compared to this half-map (FSCwork), as well as to the other (FSCtest), to confirm the absence of overfitting (**Figure 1 - figure supplement 2**). Further details of data processing and model refinement and validation are given in **Supplementary Tables 1-24**.

---

**Figure 1 - figure supplement 1:**
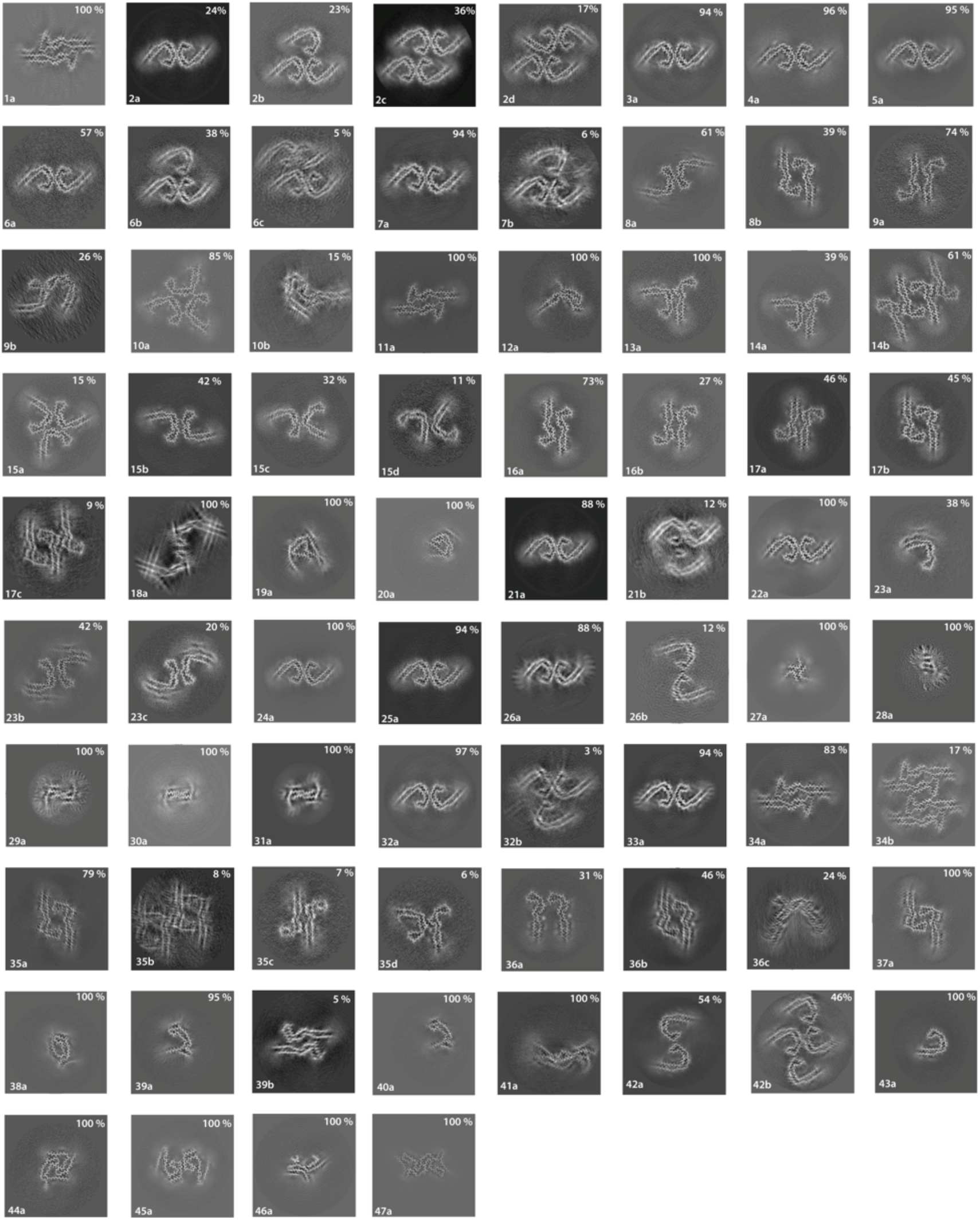
Cross-sections of cryo-EM reconstructions. Projected slices perpendicular to the helical axis, and with a thickness of approximately 4.7Å, are shown for all cryo-EM structures described in this paper. For each structure, the filament type (as defined in Table 1) is displayed in the bottom left, and the percentage of a given type in the cryo-EM data set is shown in the top right.

**Figure 1 - figure supplement 2:**
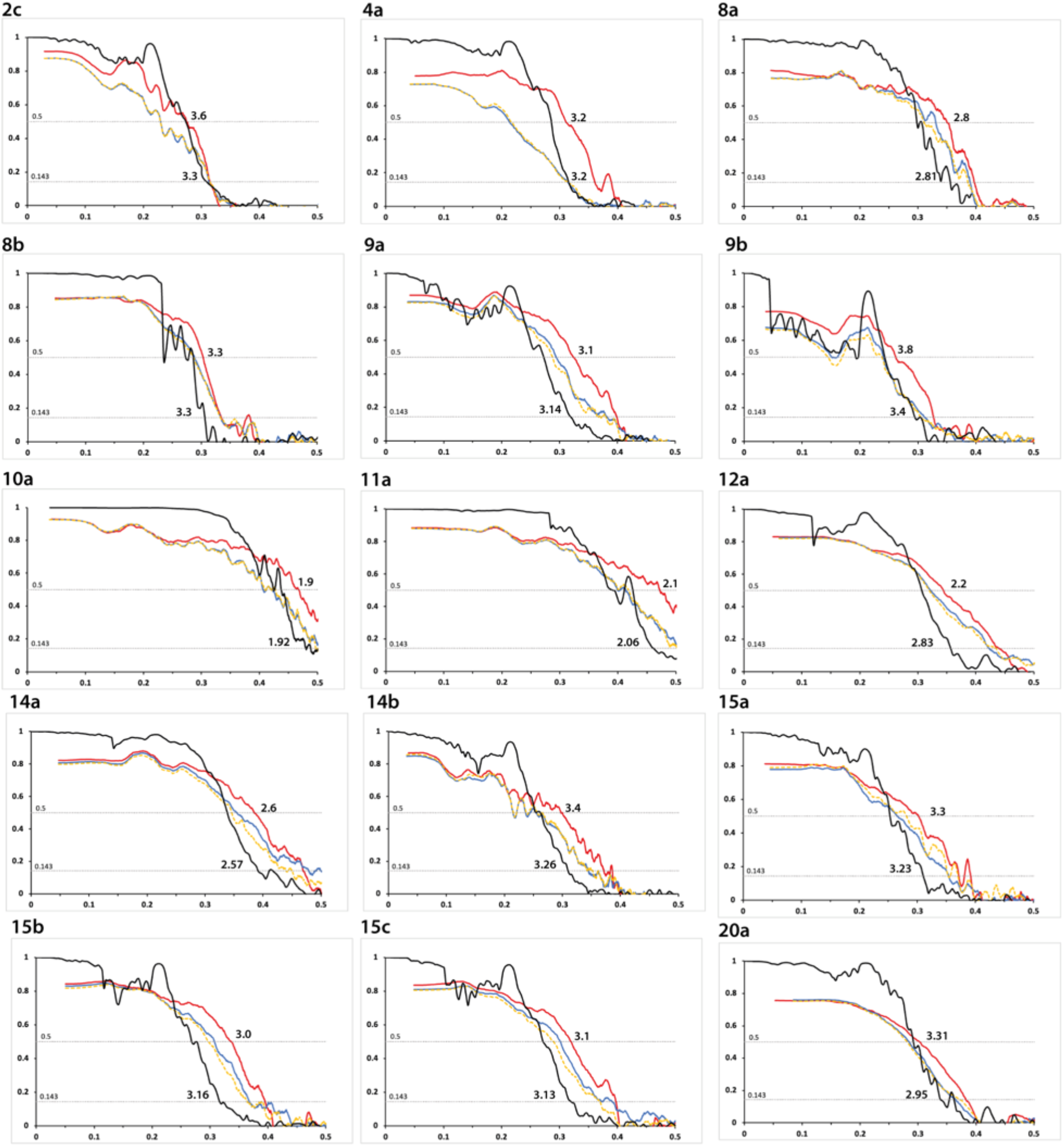
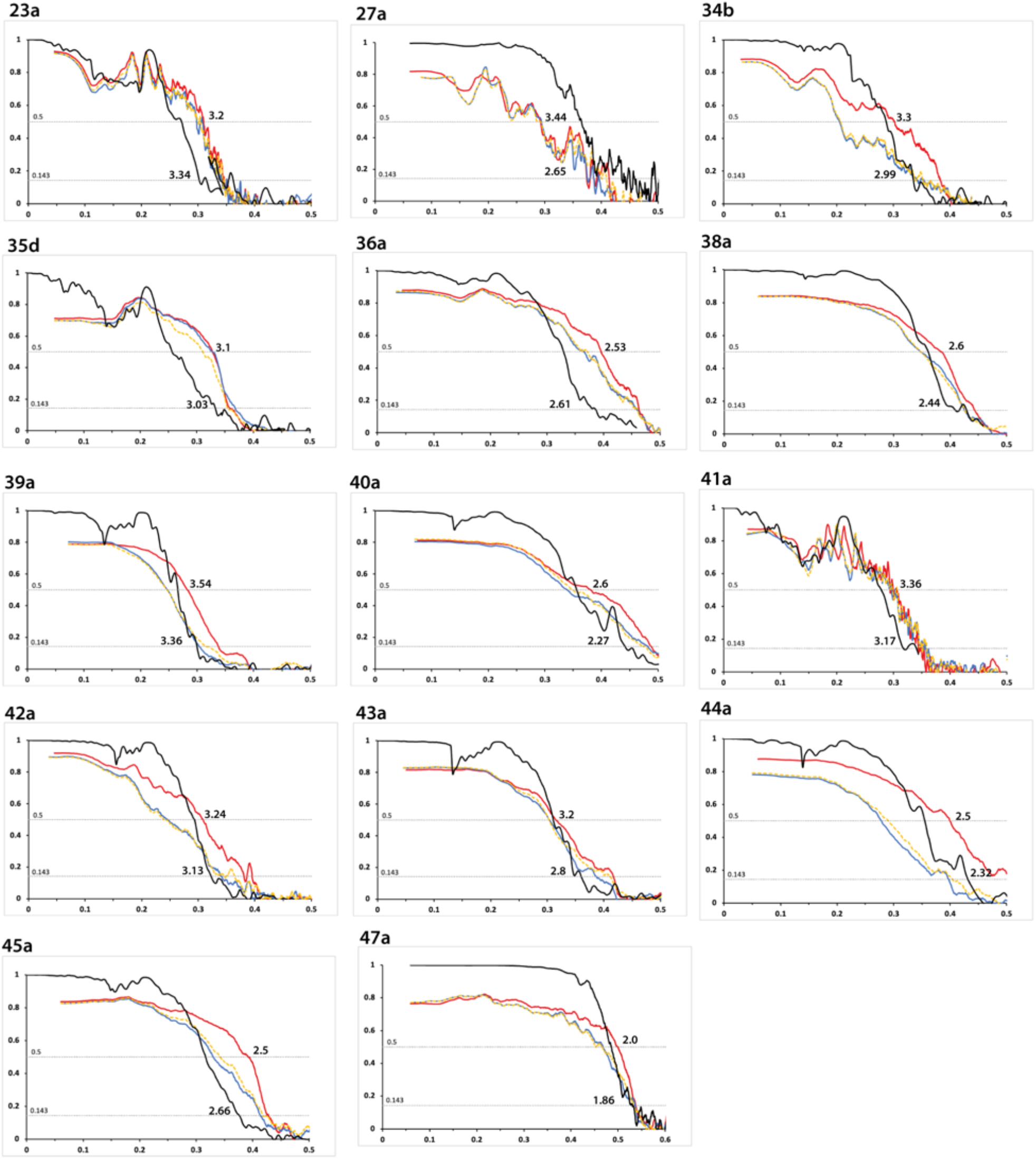
Fourier Shell Correlation (FSC) curves. For each structure described in this paper, FSC curves are shown for two independently refined cryo-EM half maps (black); for the final refined atomic model against the final cryo-EM map (red); for the atomic model refined in the first half-map against that half-map (blue); and for the refined atomic model in the first half-map against the second half-map (yellow). The resolutions where the black line drops below 0.143 and the red line drops below 0.5 are indicated. The corresponding filament type (as defined in Table 1) is shown at the top left of each graph. See Supplementary Tables 1-24 for further details on data processing.

**Figure 2 - figure supplement 1:**
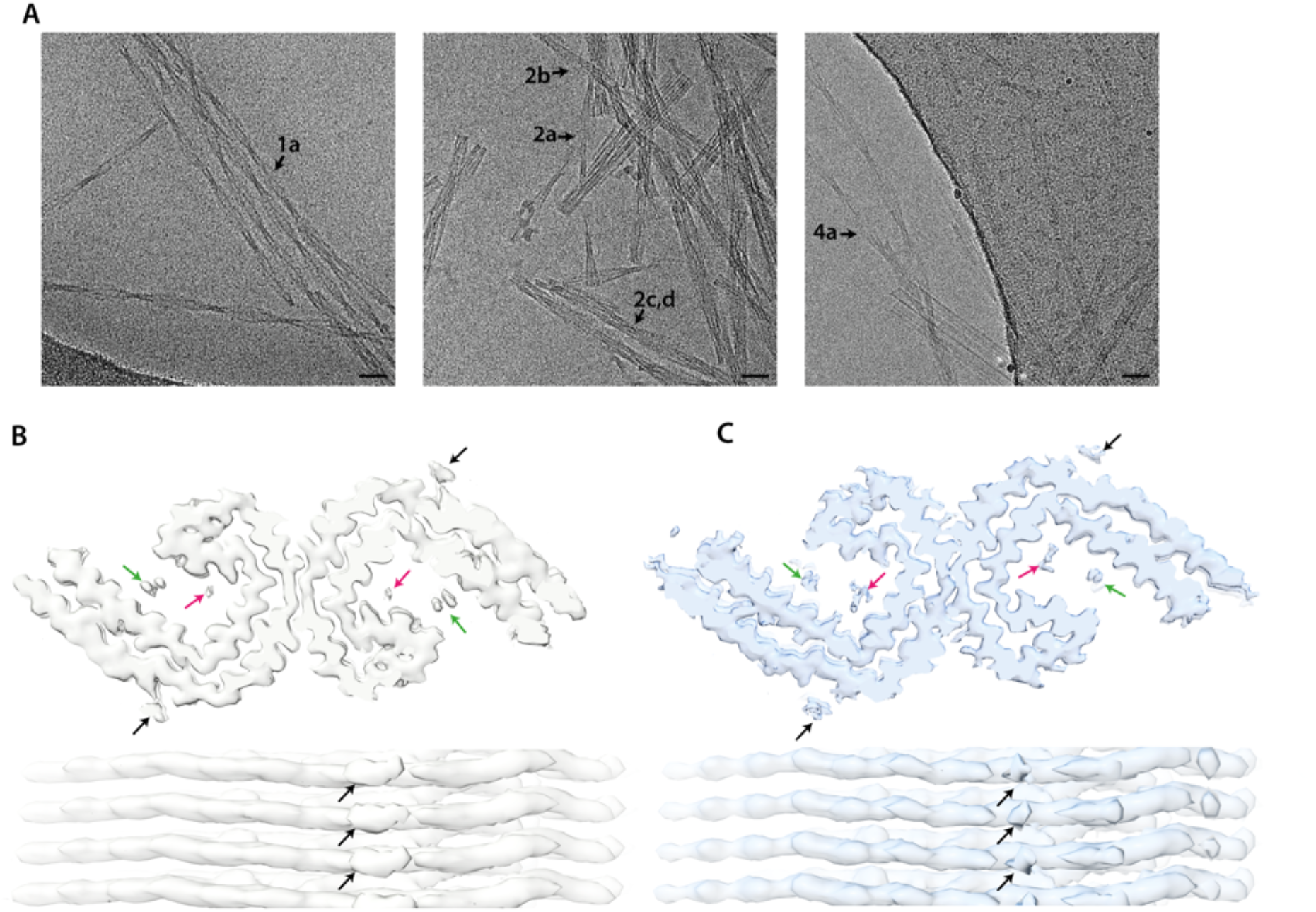
Cryo-EM micrographs and density maps comparing *in vitro* assembled PHFs and AD PHFs. **A**. Cryo-EM micrographs with filaments from *in vitro* assembly conditions 1 (left), 2 (middle) and 4 (right). Scale bar represents 100 Å. **B**. Cryo-EM density maps of *in vitro* PHF (filament type 4a, EMDB:XXXX) as top view (top) and side view (bottom). **C**. Cryo-EM density maps of AD PHF (EMDB:0259) as top view (top) and side view (bottom). The black arrows point to densities in front of lysines 317 and 321; the green and pink arrows point to densities inside the C-shape.

**Figure 3 - figure supplement 1:**
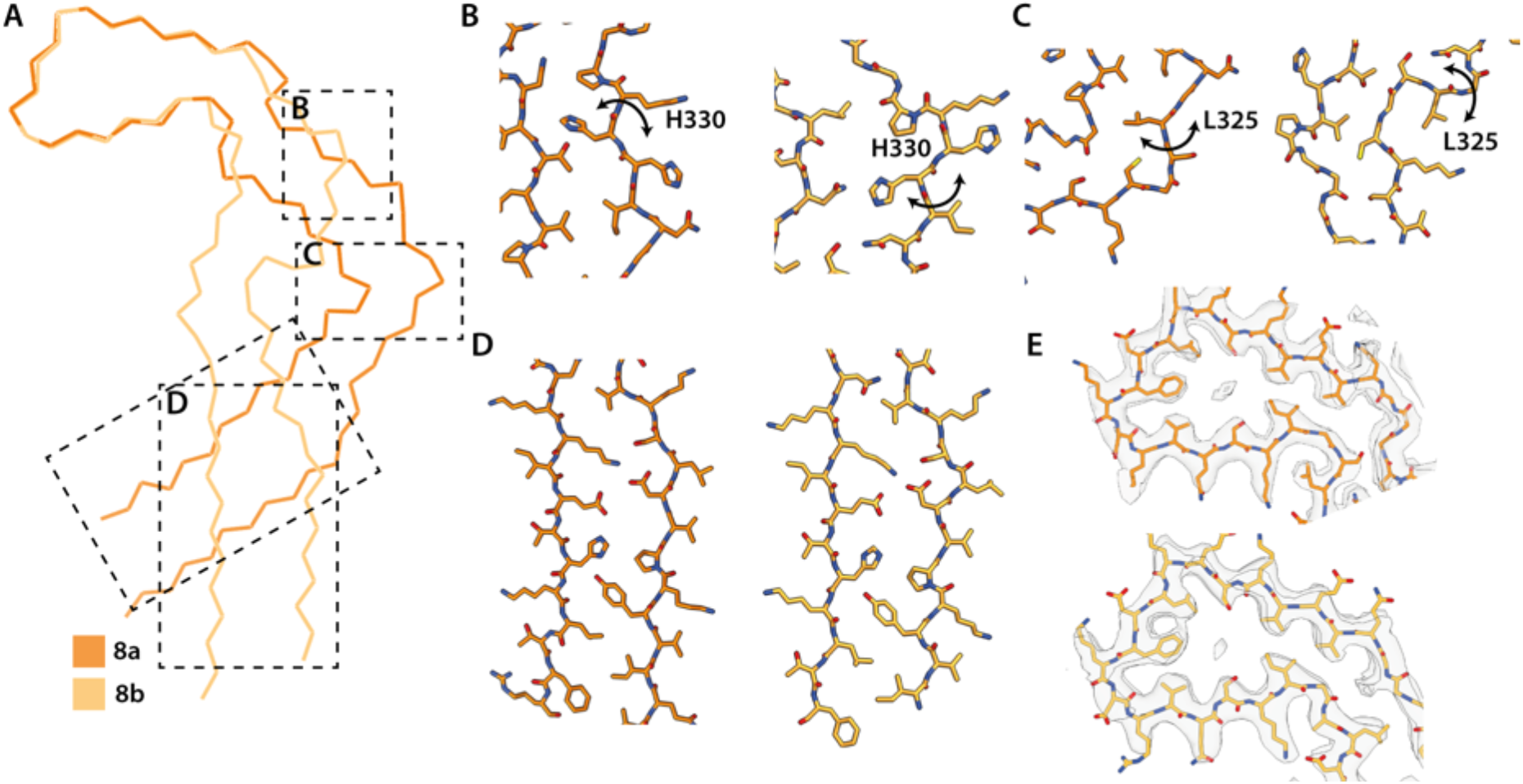
Extended and C-shaped NaCl protofilaments. **A**. Backbone ribbon view of C-shaped (filament type 8a) and extended (filament type 8b) protofilaments formed with NaCl, aligned at residues 338-354. **B-D**. Close up atomic view of regions highlighted in A. **E**. Cryo-EM density map (transparent grey) and the atomic model at residues 338-354. Filament types 8a and 8b are shown in dark and light orange, respectively.

**Figure 3 - figure supplement 2:**
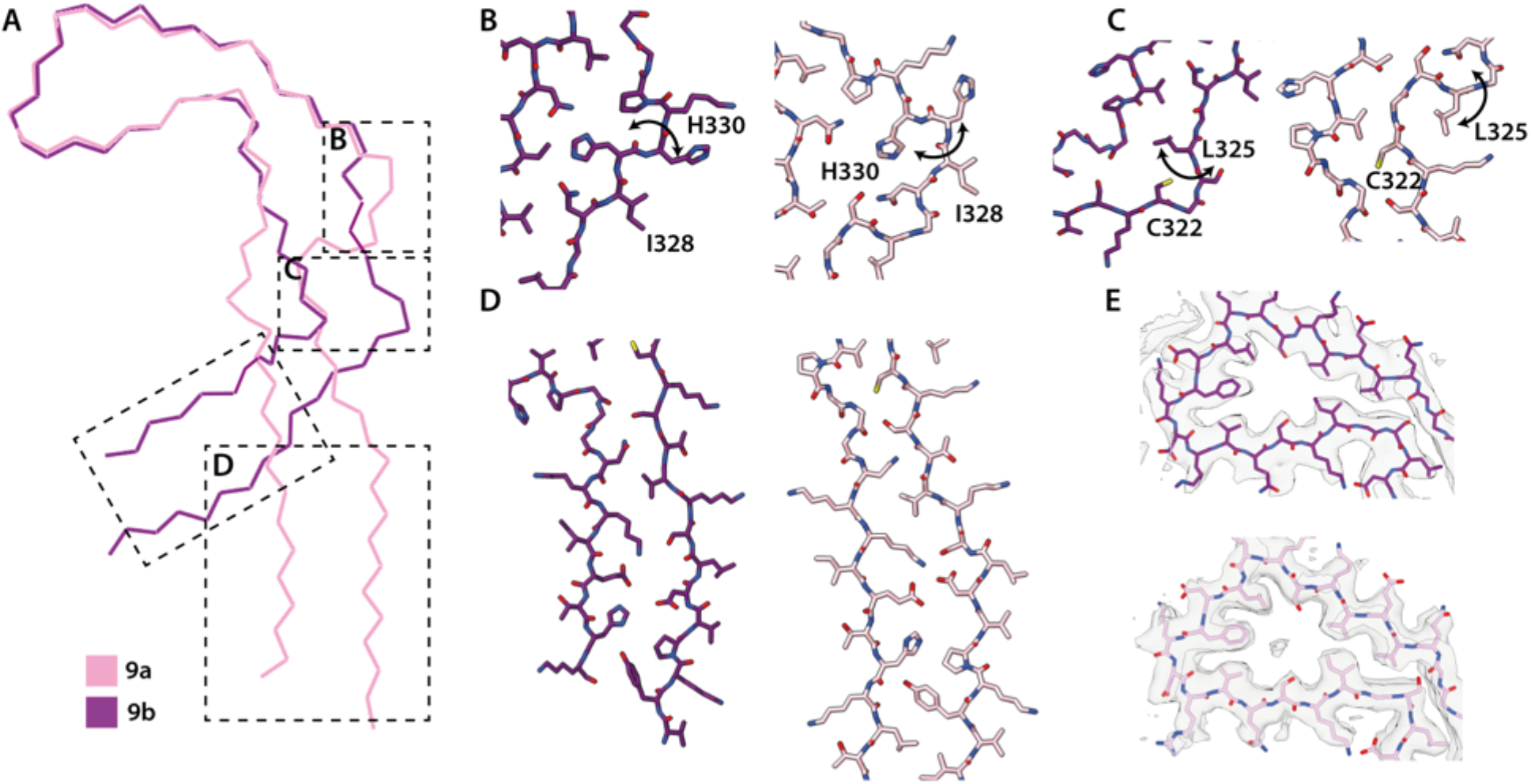
Extended and C-shaped LiCl protofilaments. **A**. Backbone ribbon view of extended (filament type 9a) and C-shaped (filament type 9b) protofilaments formed with LiCl, aligned at residues 338-354. **B-D**. Close up atomic view of regions highlighted in A. **E**. Cryo-EM density map (transparent grey) and the atomic model at residues 338-354. Filament type 9a and 9b are shown in light and dark purple, respectively.

**Figure 3 - figure supplement 3:**
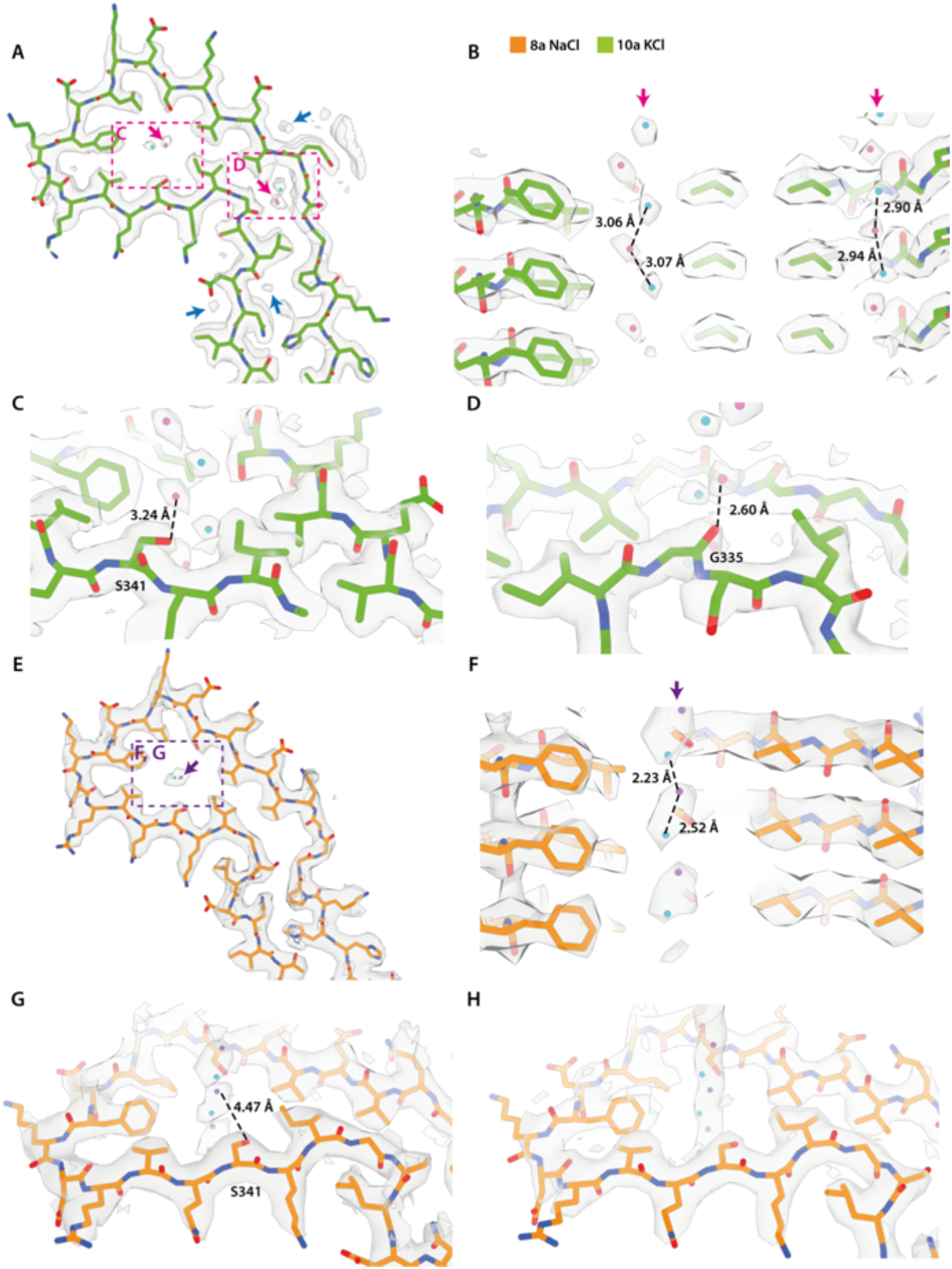
Cryo-EM densities inside the cavities of KCl and NaCl filaments. **A-H**. Cryo-EM density map (transparent grey) and the corresponding atomic models for filament types 7a (assembled with NaCl) and 8a (assembled with KCl) are shown in orange and green, respectively. **A**. Cryo-EM density at the tip of the protofilament of 8a with additional non-proteinaceous densities highlighted by arrows. Blue arrows represent putative water molecules; pink arrows represent putative K^+^ or Cl^-^ ions. K^+^ ions are shown in pink; Cl^-^ ions are shown in cyan. **B-D**. Close up view of putative K^+^ and Cl^-^ ion pairs. Distances between K^+^ and Cl^-^ ions, and Oγ of S341 or the carbonyl oxygen of G335 are indicated. **E**. Cryo-EM density of the tip of the protofilament of 8a with putative Na^+^ and Cl^-^ ions fitted into the density. Purple arrow represents putative Na^+^ or Cl^-^ ions; Na^+^ ions are shown in purple; Cl^-^ ions are shown in cyan. **F-G**. Close up view of putative Na^+^ and Cl^-^ ions. Distances between the Na^+^ and Cl^-^ ions and the Oγ of S341 are indicated. **H**. Cryo-EM density of CTE type I (EMD:0527), with putative Na^+^ and Cl^-^ ions modelled inside the additional density.

**Figure 4 - figure supplement 1:**
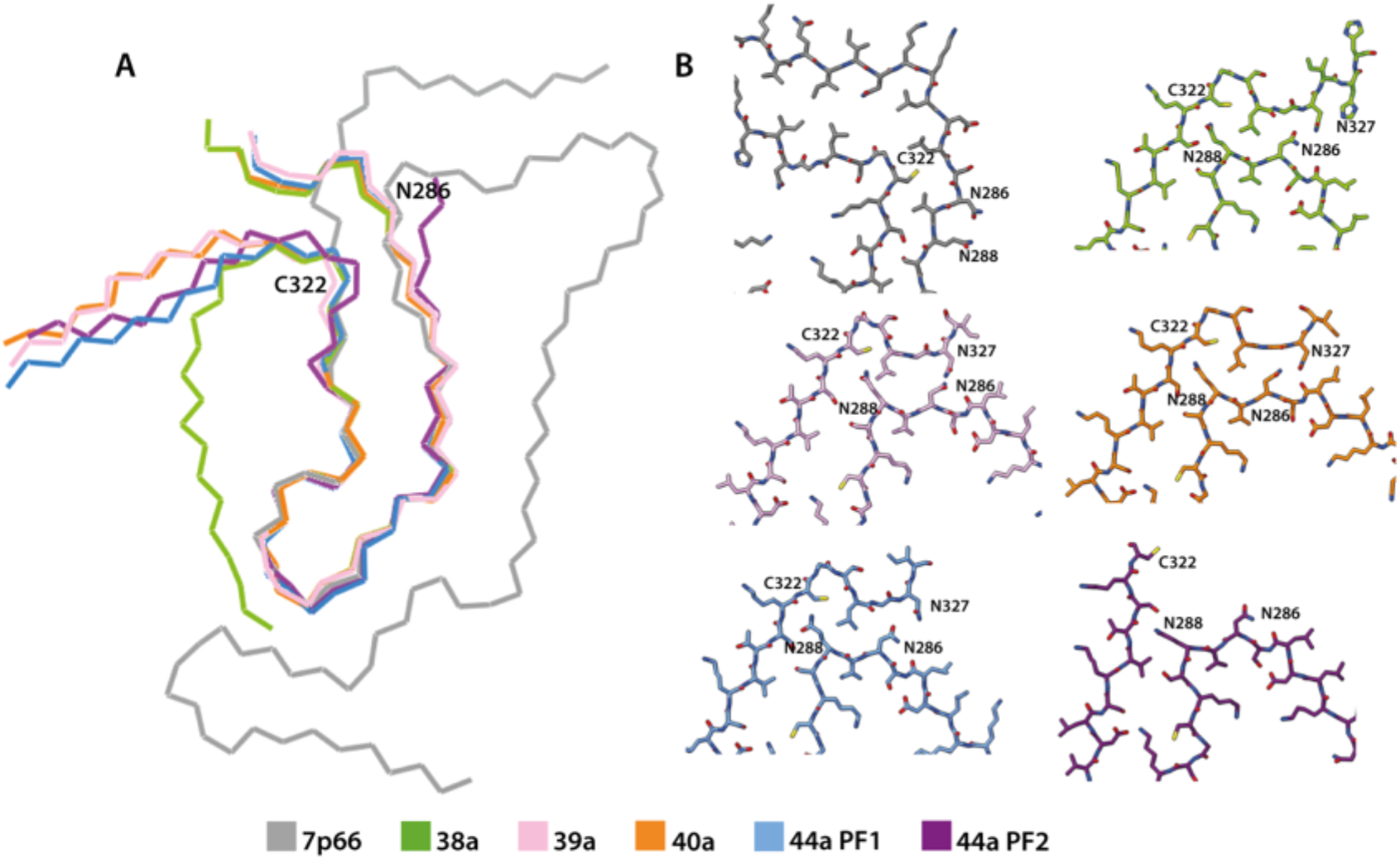
Comparison of *in vitro* assembled tau filaments with the GGT fold. **A**. Backbone ribbon view of GGT type 1 (PDB:7p66) and tau filaments from *in vitro* assembled filament types 27a, 28a, 29a, 32a protofilament (PF)1 and 32a PF2, aligned at residues 288-322. **B**. Close up atomic view. 7p66 is shown in grey, 27a in green; 28a in pink; 29a in orange; 32a PF-1 in blue and 32a PF-2 in purple.

## Cryo-EM data processing, refinement and validation statistics

**Supplementary table 1.**

| LMB Krios G2 | 2c<br>(EMDB-xxxx)<br>(PDB xxxx) |
| --- | --- |
| **Data processing** |  |
| Initial particle images (no.) | 314921 |
| Final particle images (no.) | 37499 |
| Helical twist (°) | -0.797 |
| Helical rise (Å) | 4.75 |
| Symmetry imposed | 1 |
| Map resolution FSC 0.143 (Å) | 3.29 |
| **Refinement** |  |
| Initial model used (PDB code) | 6HRE |
| Model resolution FSC 0.5 (Å) | 3.6 |
| Map sharpening *B* factor (Å<sup>2</sup>) | -22.2 |
| Model composition |  |
| Non-hydrogen atoms | 6936 |
| Protein residues | 912 |
| Ligands | 0 |
| *B* factors (Å<sup>2</sup>) |  |
| Protein | 49.38 |
| Ligand | na |
| R.m.s. deviations |  |
| Bond lengths (Å) | 0.012 |
| Bond angles (°) | 2.254 |
| Validation |  |
| MolProbity score | 1.23 |
| Clashscore | 0.65 |
| Poor rotamers (%) | 0 |
| Ramachandran plot |  |
| Favored (%) | 90.54 |
| Allowed (%) | 9.47 |
| Disallowed (%) | 0 |

**Supplementary table 2.**

| LMB Krios G1 | 4a<br>(EMDB-xxxx)<br>(PDB xxxx) |
| --- | --- |
| **Data processing** |  |
| Initial particle images (no.) | 114 023 |
| Final particle images (no.) | 40521 |
| Helical twist (°) | 179.47 |
| Helical rise (Å) | 2.358 |
| Symmetry imposed | 2<sub>1</sub> |
| Map resolution FSC 0.143 (Å) | 3.20 |
| **Refinement** |  |
| Initial model used (PDB code) | na |
| Model resolution FSC 0.5 (Å) | 3.2 |
| Map sharpening *B* factor (Å<sup>2</sup>) | -62.41 |
| Model composition |  |
| Non-hydrogen atoms | 3468 |
| Protein residues | 456 |
| Ligands |  |
| *B* factors (Å<sup>2</sup>) |  |
| Protein | 50.01 |
| Ligand | NA |
| R.m.s. deviations |  |
| Bond lengths (Å) | 0.011 |
| Bond angles (°) | 1.993 |
| Validation |  |
| MolProbity score | 1.1 |
| Clashscore | 0.28 |
| Poor rotamers (%) | 0 |
| Ramachandran plot |  |
| Favored (%) | 91.67 |
| Allowed (%) | 8.33 |
| Disallowed (%) | 0 |

**Supplementary table 3.**
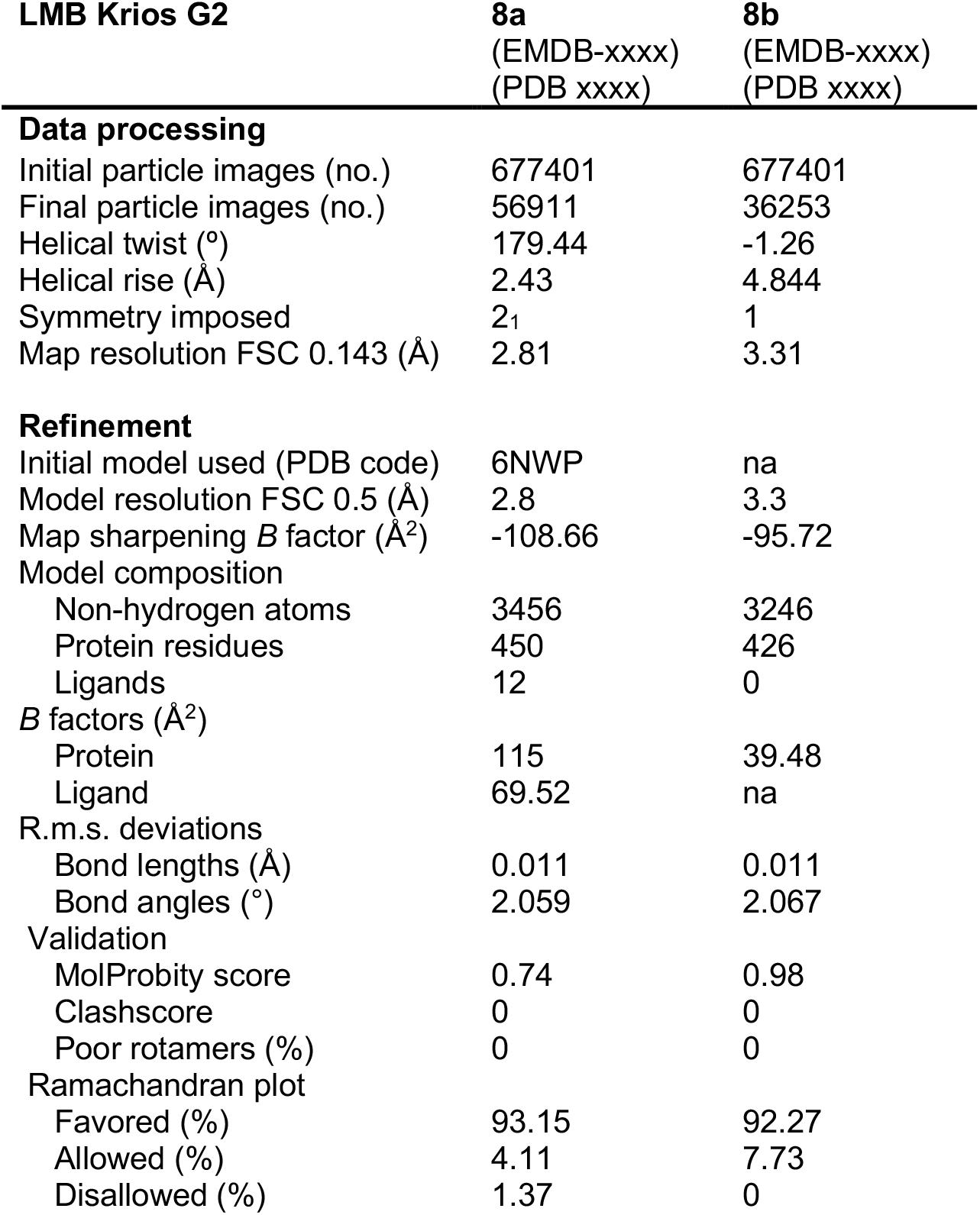

| **LMB Krios G2** | **8a**<br>(EMDB-xxxx)<br>(PDB xxxx) | **8b**<br>(EMDB-xxxx)<br>(PDB xxxx) |
| --- | --- | --- |
| **Data processing** |  |  |
| Initial particle images (no.) | 677401 | 677401 |
| Final particle images (no.) | 56911 | 36253 |
| Helical twist (°) | 179.44 | -1.26 |
| Helical rise (Å) | 2.43 | 4.844 |
| Symmetry imposed | 2<sub>1</sub> | 1 |
| Map resolution FSC 0.143 (Å) | 2.81 | 3.31 |
| **Refinement** |  |  |
| Initial model used (PDB code) | 6NWP | na |
| Model resolution FSC 0.5 (Å) | 2.8 | 3.3 |
| Map sharpening *B* factor (Å<sup>2</sup>) | -108.66 | -95.72 |
| Model composition |  |  |
| Non-hydrogen atoms | 3456 | 3246 |
| Protein residues | 450 | 426 |
| Ligands | 12 | 0 |
| *B* factors (Å<sup>2</sup>) |  |  |
| Protein | 115 | 39.48 |
| Ligand | 69.52 | na |
| R.m.s. deviations |  |  |
| Bond lengths (Å) | 0.011 | 0.011 |
| Bond angles (°) | 2.059 | 2.067 |
| Validation |  |  |
| MolProbity score | 0.74 | 0.98 |
| Clashscore | 0 | 0 |
| Poor rotamers (%) | 0 | 0 |
| Ramachandran plot |  |  |
| Favored (%) | 93.15 | 92.27 |
| Allowed (%) | 4.11 | 7.73 |
| Disallowed (%) | 1.37 | 0 |

**Supplementary table 4.**

| **TFS Krios G4** | **9a**<br>(EMDB-xxxx)<br>(PDB xxxx) | **9b**<br>(EMDB-xxxx)<br>(PDB xxxx) |
| --- | --- | --- |
| **Data processing** |  |  |
| Initial particle images (no.) | 91908 | 91908 |
| Final particle images (no.) | 24754 | 8701 |
| Helical twist (°) | -1.26 | -1.02 |
| Helical rise (Å) | 4.74 | 4.74 |
| Symmetry imposed | 2 | 1 |
| Map resolution FSC 0.143 (Å) | 3.14 | 3.4 |
| **Refinement** |  |  |
| Initial model used (PDB code) | na | na |
| Model resolution FSC 0.5 (Å) | 3.10 | 3.8 |
| Map sharpening *B* factor (Å<sup>2</sup>) | -32.69 | -14.56 |
| Model composition |  |  |
| Non-hydrogen atoms | 3444 | 3378 |
| Protein residues | 450 | 444 |
| Ligands | 0 | 0 |
| *B* factors (Å<sup>2</sup>) |  |  |
| Protein | 38.93 | 39.11 |
| Ligand | na | na |
| R.m.s. deviations |  |  |
| Bond lengths (Å) | 0.010 | 0.011 |
| Bond angles (°) | 1.965 | 2.13 |
| Validation |  |  |
| MolProbity score | 0.89 | 0.99 |
| Clashscore | 0 | 0.24 |
| Poor rotamers (%) | 0.25 | 0 |
| Ramachandran plot |  |  |
| Favored (%) | 94.29 | 94.44 |
| Allowed (%) | 5.71 | 5.32 |
| Disallowed (%) | 0 | 0.23 |

**Supplementary table 5.**

| LMB Krios G2 | 10a<br>(EMDB-xxxx)<br>(PDB xxxx) |
| --- | --- |
| **Data processing** |  |
| Initial particle images (no.) | 2087042 |
| Final particle images (no.) | 108932 |
| Helical twist (°) | -1.129 |
| Helical rise (Å) | 4.75 |
| Symmetry imposed | 3 |
| Map resolution FSC 0.143 (Å) | 1.92 |
| **Refinement** |  |
| Initial model used (PDB code) | na |
| Model resolution FSC 0.5 (Å) | 1.90 |
| Map sharpening *B* factor (Å<sup>2</sup>) | -13.20 |
| Model composition |  |
| Non-hydrogen atoms | 5112 |
| Protein residues | 666 |
| Ligands | 0 |
| *B* factors (Å<sup>2</sup>) |  |
| Protein | 49.46 |
| Ligand | na |
| R.m.s. deviations |  |
| Bond lengths (Å) | 0.011 |
| Bond angles (°) | 1.977 |
| Validation |  |
| MolProbity score | 0.81 |
| Clashscore | 0 |
| Poor rotamers (%) | 0 |
| Ramachandran plot |  |
| Favored (%) | 95.52 |
| Allowed (%) | 4.48 |
| Disallowed (%) | 0 |

**Supplementary table 6.**
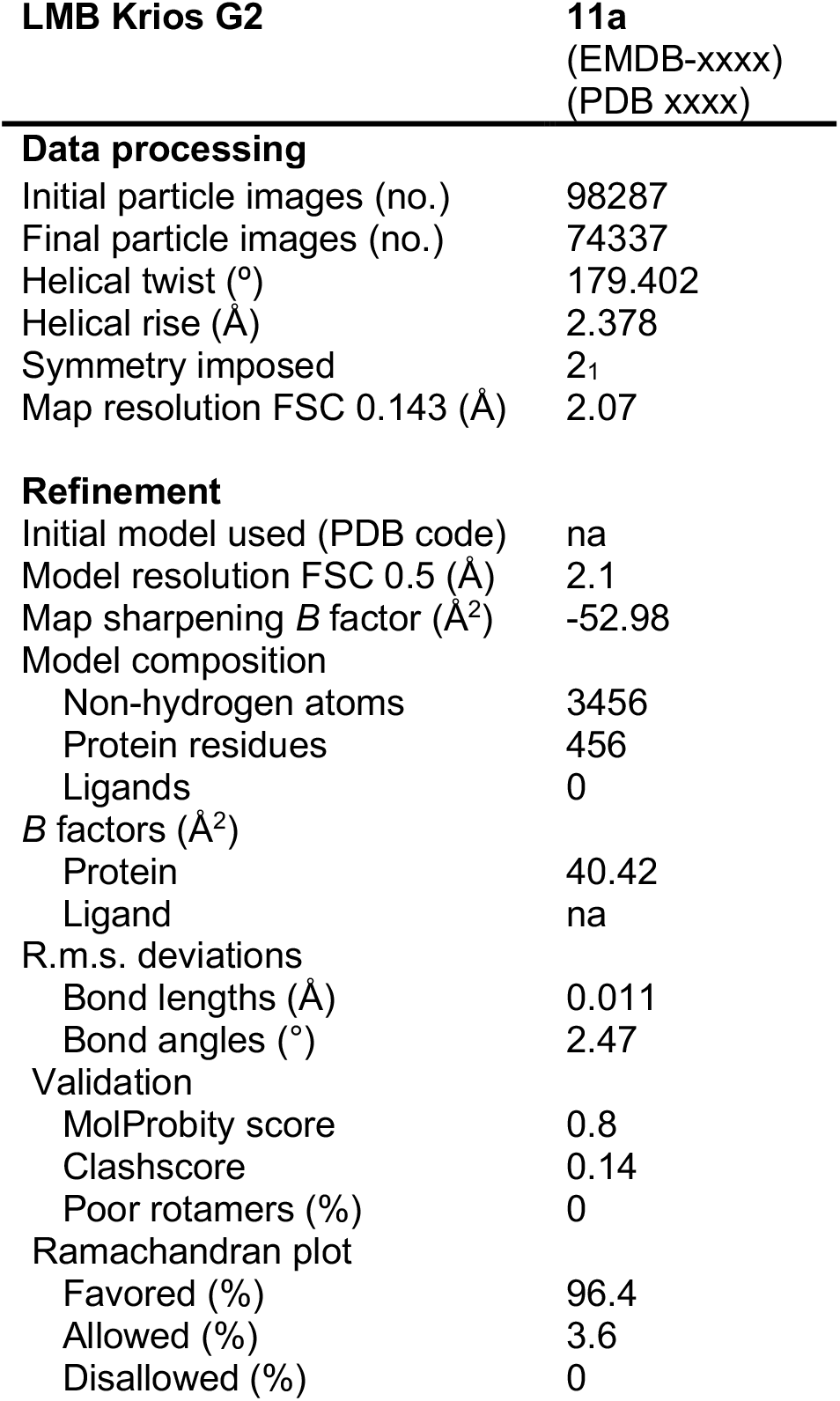

**Supplementary table 7.**

| LMB Krios G2 | 12a<br>(EMDB-xxxx)<br>(PDB xxxx) |
| --- | --- |
| **Data processing** |  |
| Initial particle images (no.) | 51533 |
| Final particle images (no.) | 17783 |
| Helical twist (°) | -1.69 |
| Helical rise (Å) | 4.8 |
| Symmetry imposed | 1 |
| Map resolution FSC 0.143 (Å) | 2.83 |
| **Refinement** |  |
| Initial model used (PDB code) | na |
| Model resolution FSC 0.5 (Å) | 2.2 |
| Map sharpening *B* factor (Å<sup>2</sup>) | -35.91 |
| Model composition |  |
| Non-hydrogen atoms | 1857 |
| Protein residues | 249 |
| Ligands | 0 |
| *B* factors (Å<sup>2</sup>) |  |
| Protein | 52.51 |
| Ligand | na |
| R.m.s. deviations |  |
| Bond lengths (Å) | 0.011 |
| Bond angles (°) | 2.336 |
| Validation |  |
| MolProbity score | 1.1 |
| Clashscore | 0.26 |
| Poor rotamers (%) | 0 |
| Ramachandran plot |  |
| Favored (%) | 91.56 |
| Allowed (%) | 8.44 |
| Disallowed (%) | 0 |

**Supplementary table 8.**

| **TFS Krios G4** | **14a**<br>(EMDB-xxxx)<br>(PDB xxxx) | **14b**<br>(EMDB-xxxx)<br>(PDB xxxx) |
| --- | --- | --- |
| **Data processing** |  |  |
| Initial particle images (no.) | 224815 | 224815 |
| Final particle images (no.) | 18013 | 28075 |
| Helical twist (°) | -1.25 | -0.99 |
| Helical rise (Å) | 4.75 | 4.78 |
| Symmetry imposed | 1 | 1 |
| Map resolution FSC 0.143 (Å) | 2.57 | 3.26 |
| **Refinement** |  |  |
| Initial model used (PDB code) | na | na |
| Model resolution FSC 0.5 (Å) | 2.6 | 3.4 |
| Map sharpening *B* factor (Å<sup>2</sup>) | -29.2 | -66.5 |
| Model composition |  |  |
| Non-hydrogen atoms | 3456 | 6888 |
| Protein residues | 453 | 900 |
| Ligands | 0 | 0 |
| *B* factors (Å<sup>2</sup>) |  |  |
| Protein | 44.56 | 38.93 |
| Ligand | na | Na |
| R.m.s. deviations |  |  |
| Bond lengths (Å) | 0.01 | 0.012 |
| Bond angles (°) | 1.926 | 2.087 |
| Validation |  |  |
| MolProbity score | 0.97 | 1.13 |
| Clashscore | 0.28 | 0.21 |
| Poor rotamers (%) | 0 | 0 |
| Ramachandran plot |  |  |
| Favored (%) | 94.78 | 90.07 |
| Allowed (%) | 5.22 | 9.93 |
| Disallowed (%) | 0 | 0 |

**Supplementary table 9.**

| **LMB Krios G2** | **15a**<br>(EMDB-xxxx)<br>(PDB xxxx) | **15b**<br>(EMDB-xxxx)<br>(PDB xxxx) | **15c**<br>(EMDB-xxxx)<br>(PDB xxxx) |
| --- | --- | --- | --- |
| **Data processing** |  |  |  |
| Initial particle images (no.) | 122945 | 122945 | 122945 |
| Final particle images (no.) | 7127 | 19606 | 14590 |
| Helical twist (°) | -1.3 | 179.426 | -1.33 |
| Helical rise (Å) | 4.75 | 2.37 | 4.75 |
| Symmetry imposed | 3 | 2<sub>1</sub> | 1 |
| Map resolution FSC 0.143 (Å) | 3.23 | 3.16 | 3.13 |
| **Refinement** |  |  |  |
| Initial model used (PDB code) | na | na | na |
| Model resolution FSC 0.5 (Å) | 3.3 | 3.0 | 3.1 |
| Map sharpening *B* factor (Å<sup>2</sup>) | -54.72 | -45.01 | -46.55 |
| Model composition |  |  |  |
| Non-hydrogen atoms | 4698 | 3342 | 3132 |
| Protein residues | 621 | 438 | 414 |
| Ligands | 0 | 0 | 0 |
| *B* factors (Å<sup>2</sup>) |  |  |  |
| Protein | 101.29 | 110.93 | 101.29 |
| Ligand | na | na | na |
| R.m.s. deviations |  |  |  |
| Bond lengths (Å) | 0.011 | 0.01 | 0.01 |
| Bond angles (°) | 1.97 | 1.88 | 1.948 |
| Validation |  |  |  |
| MolProbity score | 0.93 | 0.71 | 0.77 |
| Clashscore | 0.63 | 0 | 0 |
| Poor rotamers (%) | 0 | 0 | 0 |
| Ramachandran plot |  |  |  |
| Favored (%) | 96.52 | 96.71 | 96.02 |
| Allowed (%) | 3.48 | 3.29 | 3.98 |
| Disallowed (%) | 0 | 0 | 0 |

**Supplementary table 10.**
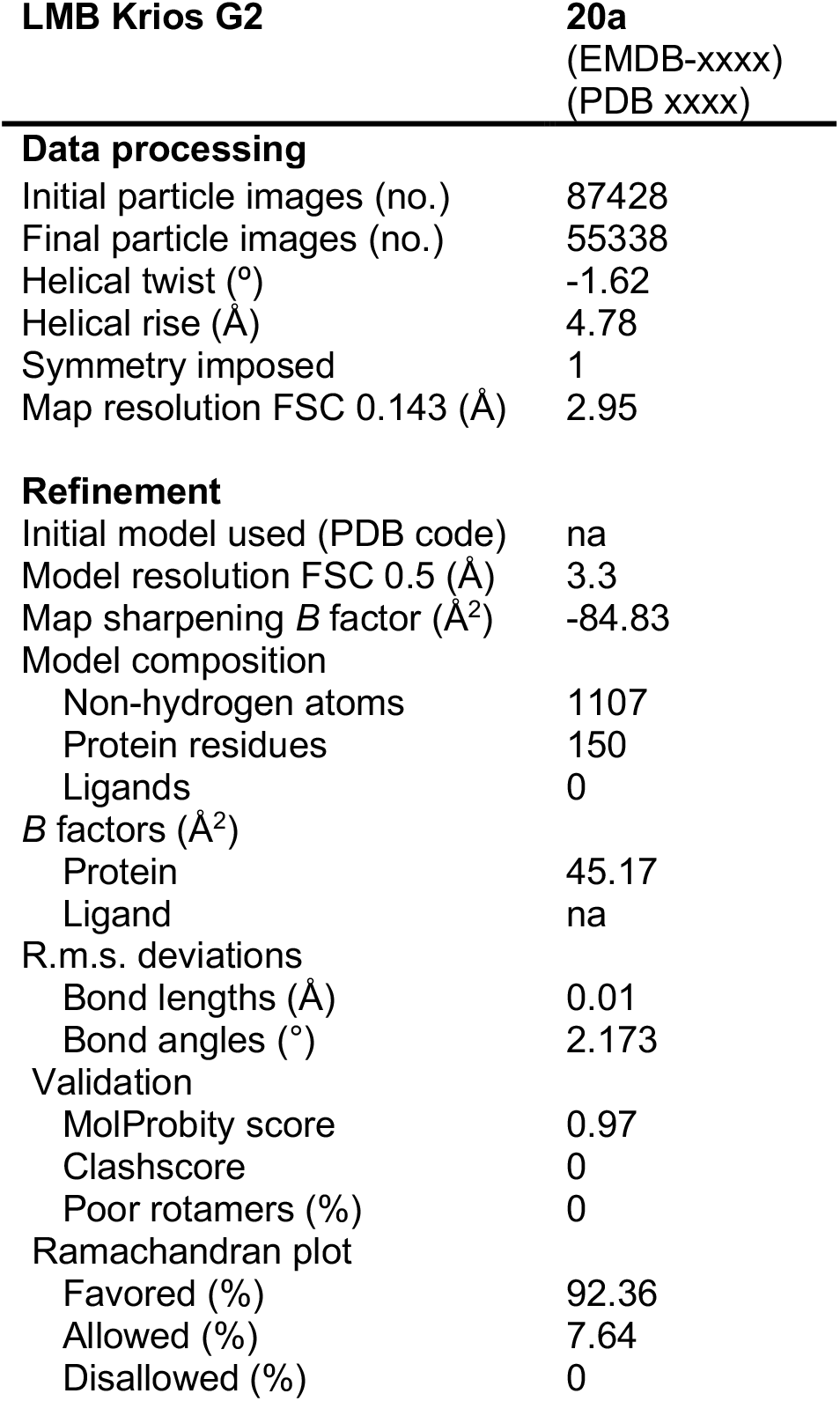

**Supplementary table 11.**

| TFS Krios G4 | 23a<br>(EMDB-xxxx)<br>(PDB xxxx) |
| --- | --- |
| **Data processing** |  |
| Initial particle images (no.) | 147647 |
| Final particle images (no.) | 23243 |
| Helical twist (°) | -1.25 |
| Helical rise (Å) | 4.75 |
| Symmetry imposed | 1 |
| Map resolution FSC 0.143 (Å) | 3.34 |
| **Refinement** |  |
| Initial model used (PDB code) | na |
| Model resolution FSC 0.5 (Å) | 3.2 |
| Map sharpening *B* factor (Å<sup>2</sup>) | -55.26 |
| Model composition |  |
| Non-hydrogen atoms | 1107 |
| Protein residues | 150 |
| Ligands | 0 |
| *B* factors (Å<sup>2</sup>) |  |
| Protein | 46.71 |
| Ligand | na |
| R.m.s. deviations |  |
| Bond lengths (Å) | 0.01 |
| Bond angles (°) | 2.061 |
| Validation |  |
| MolProbity score | 1.08 |
| Clashscore | 0.29 |
| Poor rotamers (%) | 0 |
| Ramachandran plot |  |
| Favored (%) | 92.49 |
| Allowed (%) | 7.51 |
| Disallowed (%) | 0 |

**Supplementary table 12.**

| TFS Glacios | 27a<br>(EMDB-xxxx)<br>(PDB xxxx) |
| --- | --- |
| **Data processing** |  |
| Initial particle images (no.) | 1005768 |
| Final particle images (no.) | 54983 |
| Helical twist (°) | -0.75 |
| Helical rise (Å) | 4.70 |
| Symmetry imposed | 3 |
| Map resolution FSC 0.143 (Å) | 2.65 |
| **Refinement** |  |
| Initial model used (PDB code) | na |
| Model resolution FSC 0.5 (Å) | 3.44 |
| Map sharpening *B* factor (Å<sup>2</sup>) | -44.56 |
| Model composition |  |
| Non-hydrogen atoms | 810 |
| Protein residues | 99 |
| Ligands | 0 |
| *B* factors (Å<sup>2</sup>) |  |
| Protein | 52.47 |
| Ligand | na |
| R.m.s. deviations |  |
| Bond lengths (Å) | 0.012 |
| Bond angles (°) | 2.833 |
| Validation |  |
| MolProbity score | 1.24 |
| Clashscore | 0.59 |
| Poor rotamers (%) | 0 |
| Ramachandran plot |  |
| Favored (%) | 90.12 |
| Allowed (%) | 9.88 |
| Disallowed (%) | 0 |

**Supplementary table 13.**
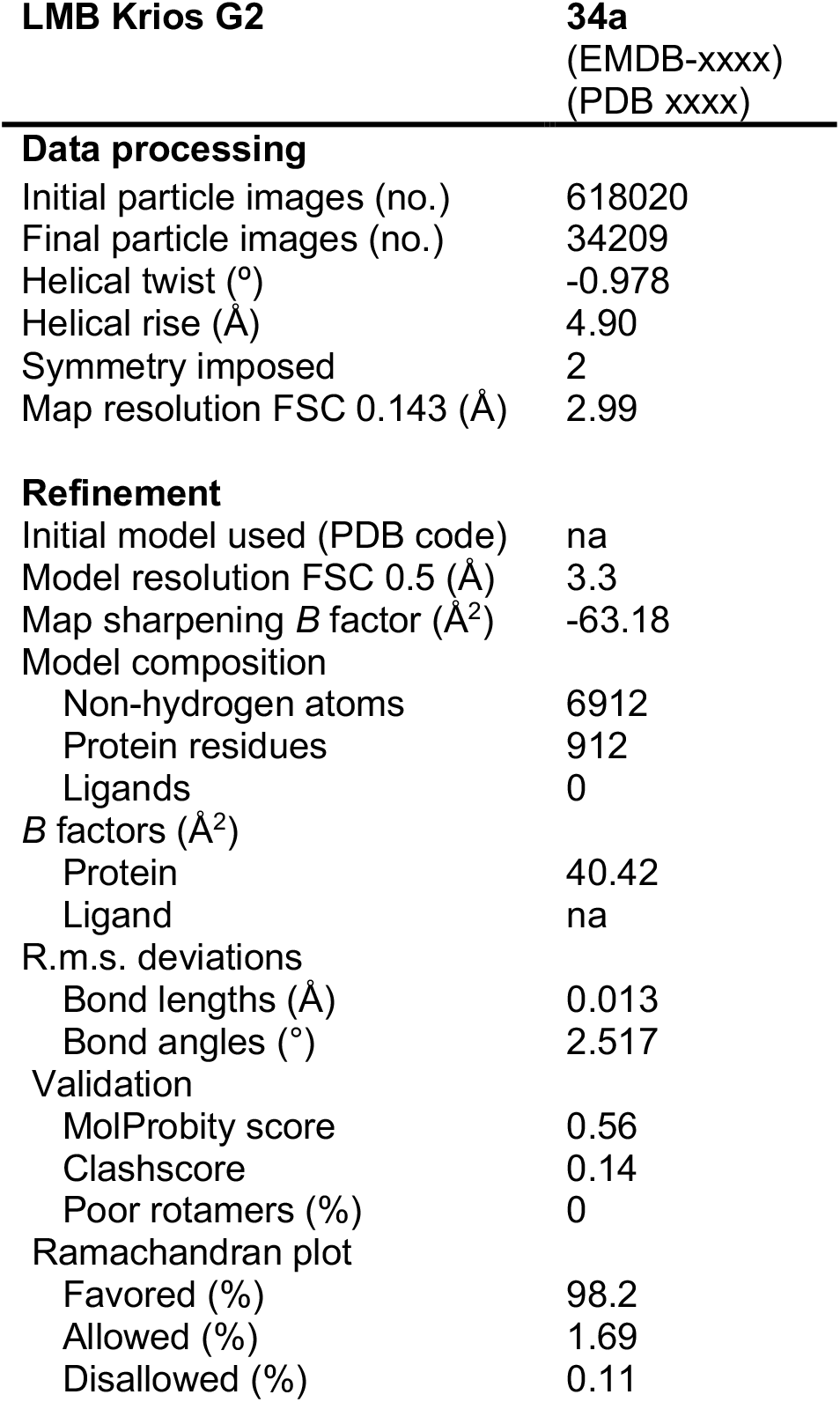

| LMB Krios G2 | 34a<br>(EMDB-xxxx)<br>(PDB xxxx) |
| --- | --- |
| **Data processing** |  |
| Initial particle images (no.) | 618020 |
| Final particle images (no.) | 34209 |
| Helical twist (°) | -0.978 |
| Helical rise (Å) | 4.90 |
| Symmetry imposed | 2 |
| Map resolution FSC 0.143 (Å) | 2.99 |
| **Refinement** |  |
| Initial model used (PDB code) | na |
| Model resolution FSC 0.5 (Å) | 3.3 |
| Map sharpening *B* factor (Å<sup>2</sup>) | -63.18 |
| Model composition |  |
| Non-hydrogen atoms | 6912 |
| Protein residues | 912 |
| Ligands | 0 |
| *B* factors (Å<sup>2</sup>) |  |
| Protein | 40.42 |
| Ligand | na |
| R.m.s. deviations |  |
| Bond lengths (Å) | 0.013 |
| Bond angles (°) | 2.517 |
| Validation |  |
| MolProbity score | 0.56 |
| Clashscore | 0.14 |
| Poor rotamers (%) | 0 |
| Ramachandran plot |  |
| Favored (%) | 98.2 |
| Allowed (%) | 1.69 |
| Disallowed (%) | 0.11 |

**Supplementary table 14.**

| TFS Krios G4 | 35d<br>(EMDB-xxxx)<br>(PDB xxxx) |
| --- | --- |
| **Data processing** |  |
| Initial particle images (no.) | 214567 |
| Final particle images (no.) | 7045 |
| Helical twist (°) | -1.25 |
| Helical rise (Å) | 4.77 |
| Symmetry imposed | 1 |
| Map resolution FSC 0.143 (Å) | 3.03 |
| **Refinement** |  |
| Initial model used (PDB code) | na |
| Model resolution FSC 0.5 (Å) | 3.1 |
| Map sharpening *B* factor (Å<sup>2</sup>) | -11.08 |
| Model composition |  |
| Non-hydrogen atoms | 3444 |
| Protein residues | 450 |
| Ligands | 0 |
| *B* factors (Å<sup>2</sup>) |  |
| Protein | 38.93 |
| Ligand | na |
| R.m.s. deviations |  |
| Bond lengths (Å) | 0.01 |
| Bond angles (°) | 2.033 |
| Validation |  |
| MolProbity score | 0.87 |
| Clashscore | 0 |
| Poor rotamers (%) | 0 |
| Ramachandran plot |  |
| Favored (%) | 94.52 |
| Allowed (%) | 5.48 |
| Disallowed (%) | 0 |

**Supplementary table 15.**

| TFS Krios G4 | 36a<br>(EMDB-xxxx)<br>(PDB xxxx) |
| --- | --- |
| **Data processing** |  |
| Initial particle images (no.) | 564821 |
| Final particle images (no.) | 31627 |
| Helical twist (°) | -1.37 |
| Helical rise (Å) | 4.74 |
| Symmetry imposed | 1 |
| Map resolution FSC 0.143 (Å) | 2.61 |
| **Refinement** |  |
| Initial model used (PDB code) | na |
| Model resolution FSC 0.5 (Å) | 2.53 |
| Map sharpening *B* factor (Å<sup>2</sup>) | -31.51 |
| Model composition |  |
| Non-hydrogen atoms | 3246 |
| Protein residues | 426 |
| Ligands | 0 |
| *B* factors (Å<sup>2</sup>) |  |
| Protein | 39.48 |
| Ligand | na |
| R.m.s. deviations |  |
| Bond lengths (Å) | 0.01 |
| Bond angles (°) | 1.969 |
| Validation |  |
| MolProbity score | 0.84 |
| Clashscore | 0.15 |
| Poor rotamers (%) | 0 |
| Ramachandran plot |  |
| Favored (%) | 95.89 |
| Allowed (%) | 4.11 |
| Disallowed (%) | 0.11 |

**Supplementary table 16.**

| TFS Krios G4 | 38a<br>(EMDB-xxxx)<br>(PDB xxxx) |
| --- | --- |
| **Data processing** |  |
| Initial particle images (no.) | 677859 |
| Final particle images (no.) | 37574 |
| Helical twist (°) | -1.49 |
| Helical rise (Å) | 4.74 |
| Symmetry imposed | 1 |
| Map resolution FSC 0.143 (Å) | 2.44 |
| **Refinement** |  |
| Initial model used (PDB code) | na |
| Model resolution FSC 0.5 (Å) | 2.61 |
| Map sharpening *B* factor (Å<sup>2</sup>) | -24.7 |
| Model composition |  |
| Non-hydrogen atoms | 1335 |
| Protein residues | 183 |
| Ligands | 0 |
| *B* factors (Å<sup>2</sup>) |  |
| Protein | 42.58 |
| Ligand | na |
| R.m.s. deviations |  |
| Bond lengths (Å) | 0.01 |
| Bond angles (°) | 2.172 |
| Validation |  |
| MolProbity score | 0.72 |
| Clashscore | 0 |
| Poor rotamers (%) | 0 |
| Ramachandran plot |  |
| Favored (%) | 96.61 |
| Allowed (%) | 3.39 |
| Disallowed (%) | 0 |

**Supplementary table 17.**
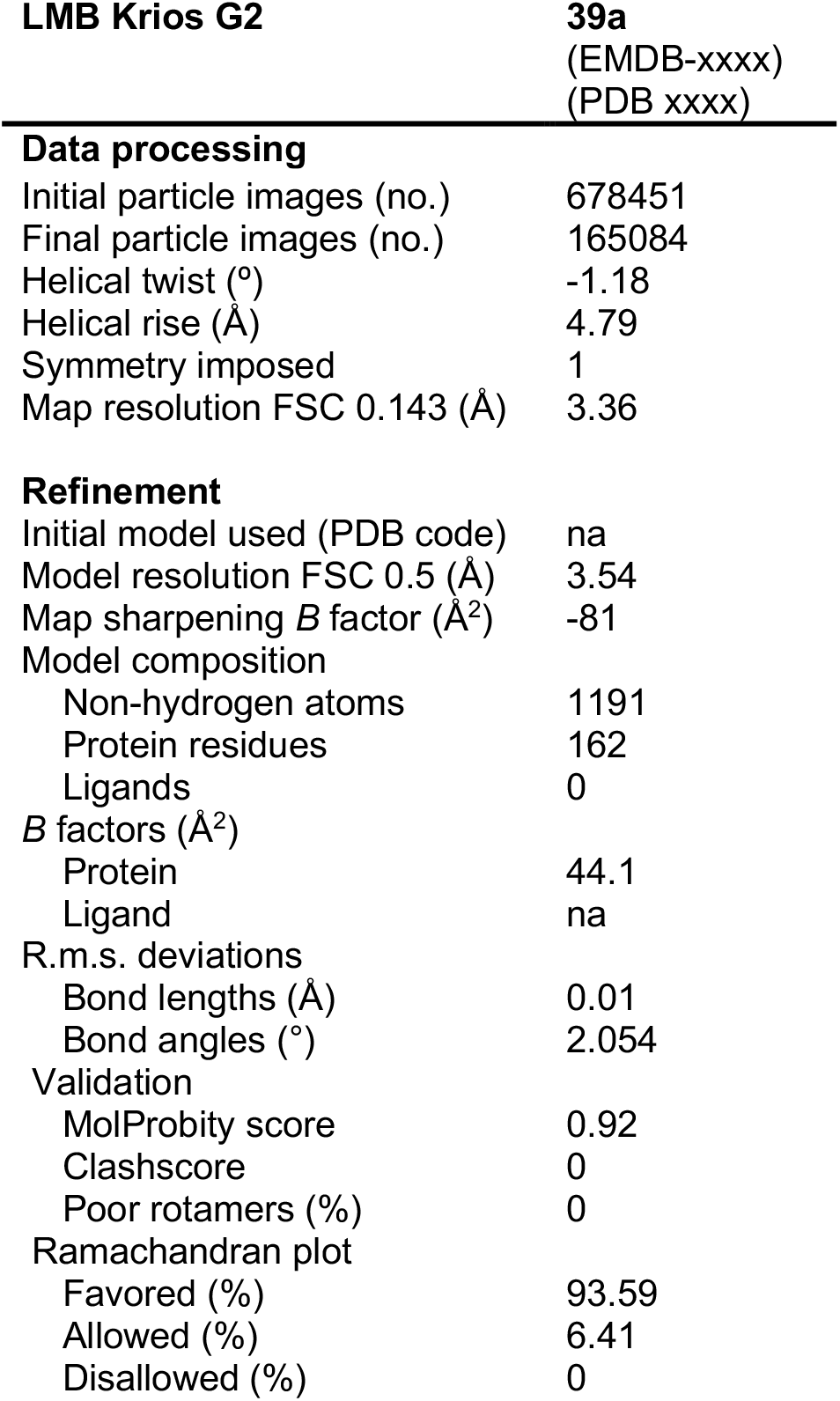

**Supplementary table 18.**
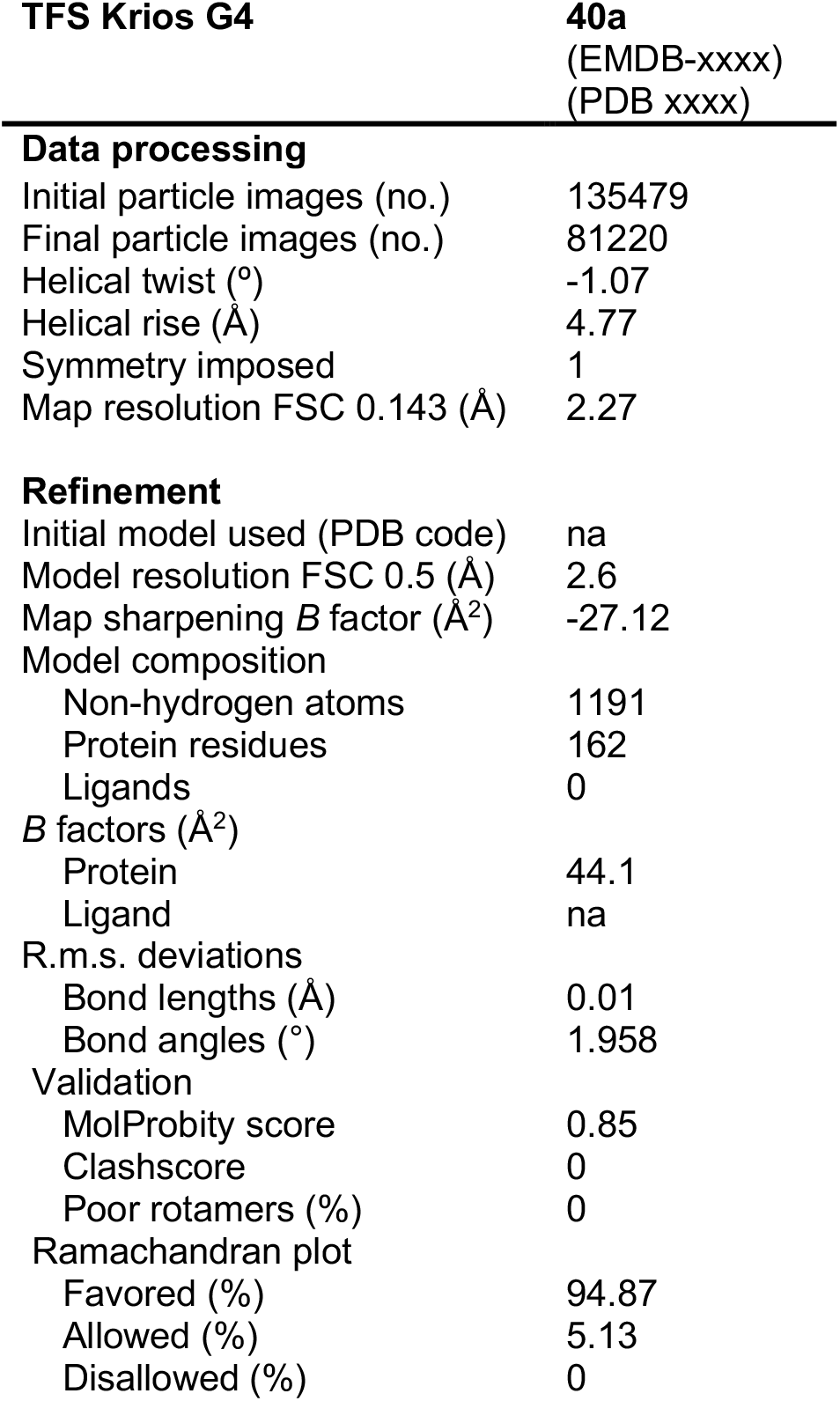

| TFS Krios G4 | 40a<br>(EMDB-xxxx)<br>(PDB xxxx) |
| --- | --- |
| **Data processing** |  |
| Initial particle images (no.) | 135479 |
| Final particle images (no.) | 81220 |
| Helical twist (°) | -1.07 |
| Helical rise (Å) | 4.77 |
| Symmetry imposed | 1 |
| Map resolution FSC 0.143 (Å) | 2.27 |
| **Refinement** |  |
| Initial model used (PDB code) | na |
| Model resolution FSC 0.5 (Å) | 2.6 |
| Map sharpening *B* factor (Å<sup>2</sup>) | -27.12 |
| Model composition |  |
| Non-hydrogen atoms | 1191 |
| Protein residues | 162 |
| Ligands | 0 |
| *B* factors (Å<sup>2</sup>) |  |
| Protein | 44.1 |
| Ligand | na |
| R.m.s. deviations |  |
| Bond lengths (Å) | 0.01 |
| Bond angles (°) | 1.958 |
| Validation |  |
| MolProbity score | 0.85 |
| Clashscore | 0 |
| Poor rotamers (%) | 0 |
| Ramachandran plot |  |
| Favored (%) | 94.87 |
| Allowed (%) | 5.13 |
| Disallowed (%) | 0 |

**Supplementary table 19.**

| TFS Krios G4 | 41a<br>(EMDB-xxxx)<br>(PDB xxxx) |
| --- | --- |
| **Data processing** |  |
| Initial particle images (no.) | 116487 |
| Final particle images (no.) | 26532 |
| Helical twist (°) | -0.807 |
| Helical rise (Å) | 4.77 |
| Symmetry imposed | 1 |
| Map resolution FSC 0.143 (Å) | 3.17 |
| **Refinement** |  |
| Initial model used (PDB code) | na |
| Model resolution FSC 0.5 (Å) | 3.36 |
| Map sharpening *B* factor (Å<sup>2</sup>) | -9.73 |
| Model composition |  |
| Non-hydrogen atoms | 1512 |
| Protein residues | 204 |
| Ligands | 0 |
| *B* factors (Å<sup>2</sup>) |  |
| Protein | 33.26 |
| Ligand | na |
| R.m.s. deviations |  |
| Bond lengths (Å) | 0.011 |
| Bond angles (°) | 2.473 |
| Validation |  |
| MolProbity score | 1.51 |
| Clashscore | 1.29 |
| Poor rotamers (%) | 0 |
| Ramachandran plot |  |
| Favored (%) | 85.42 |
| Allowed (%) | 14.58 |
| Disallowed (%) | 0 |

**Supplementary table 20.**

| LMB Krios G2 | 42a<br>(EMDB-xxxx)<br>(PDB xxxx) |
| --- | --- |
| **Data processing** |  |
| Initial particle images (no.) | 181285 |
| Final particle images (no.) | 82803 |
| Helical twist (°) | 179.59 |
| Helical rise (Å) | 2.38 |
| Symmetry imposed | 2<sub>1</sub> |
| Map resolution FSC 0.143 (Å) | 3.13 |
| **Refinement** |  |
| Initial model used (PDB code) | na |
| Model resolution FSC 0.5 (Å) | 3.24 |
| Map sharpening *B* factor (Å<sup>2</sup>) | -63.04 |
| Model composition |  |
| Non-hydrogen atoms | 3468 |
| Protein residues | 456 |
| Ligands | 0 |
| *B* factors (Å<sup>2</sup>) |  |
| Protein | 50.07 |
| Ligand | na |
| R.m.s. deviations |  |
| Bond lengths (Å) | 0.011 |
| Bond angles (°) | 2.093 |
| Validation |  |
| MolProbity score | 1.05 |
| Clashscore | 0 |
| Poor rotamers (%) | 0 |
| Ramachandran plot |  |
| Favored (%) | 89.86 |
| Allowed (%) | 10.14 |
| Disallowed (%) | 0 |

**Supplementary table 21.**

| LMB Krios G2 | 43a<br>(EMDB-xxxx)<br>(PDB xxxx) |
| --- | --- |
| **Data processing** |  |
| Initial particle images (no.) | 180999 |
| Final particle images (no.) | 84513 |
| Helical twist (°) | -1.29 |
| Helical rise (Å) | 4.77 |
| Symmetry imposed | 1 |
| Map resolution FSC 0.143 (Å) | 2.8 |
| **Refinement** |  |
| Initial model used (PDB code) | na |
| Model resolution FSC 0.5 (Å) | 3.24 |
| Map sharpening *B* factor (Å<sup>2</sup>) | -41.5 |
| Model composition |  |
| Non-hydrogen atoms | 1734 |
| Protein residues | 288 |
| Ligands | 0 |
| *B* factors (Å<sup>2</sup>) |  |
| Protein | 71.71 |
| Ligand | na |
| R.m.s. deviations |  |
| Bond lengths (Å) | 0.011 |
| Bond angles (°) | 2.261 |
| Validation |  |
| MolProbity score | 1.08 |
| Clashscore | 0 |
| Poor rotamers (%) | 0 |
| Ramachandran plot |  |
| Favored (%) | 88.74 |
| Allowed (%) | 11.26 |
| Disallowed (%) | 0 |

**Supplementary table 22.**
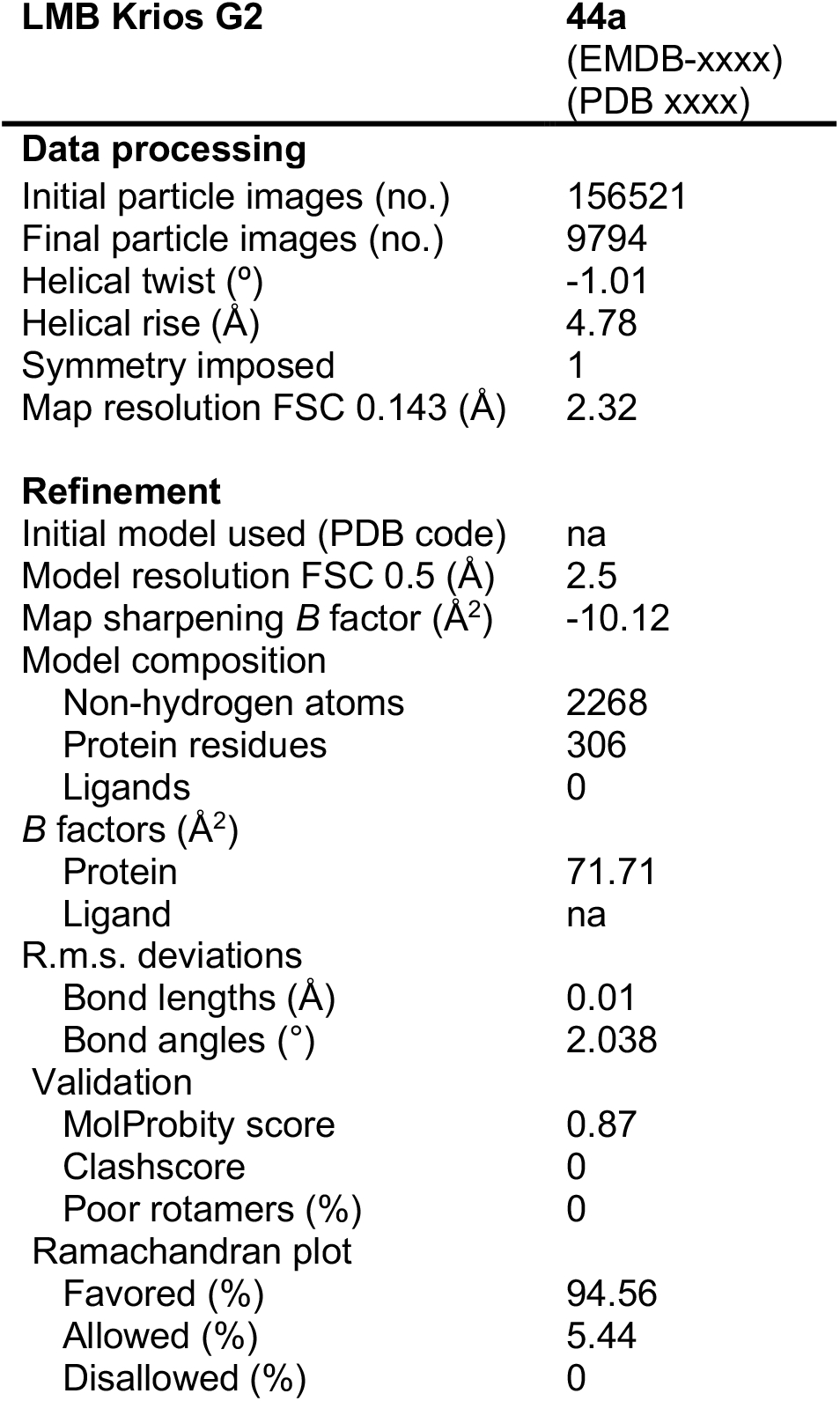

**Supplementary table 23.**
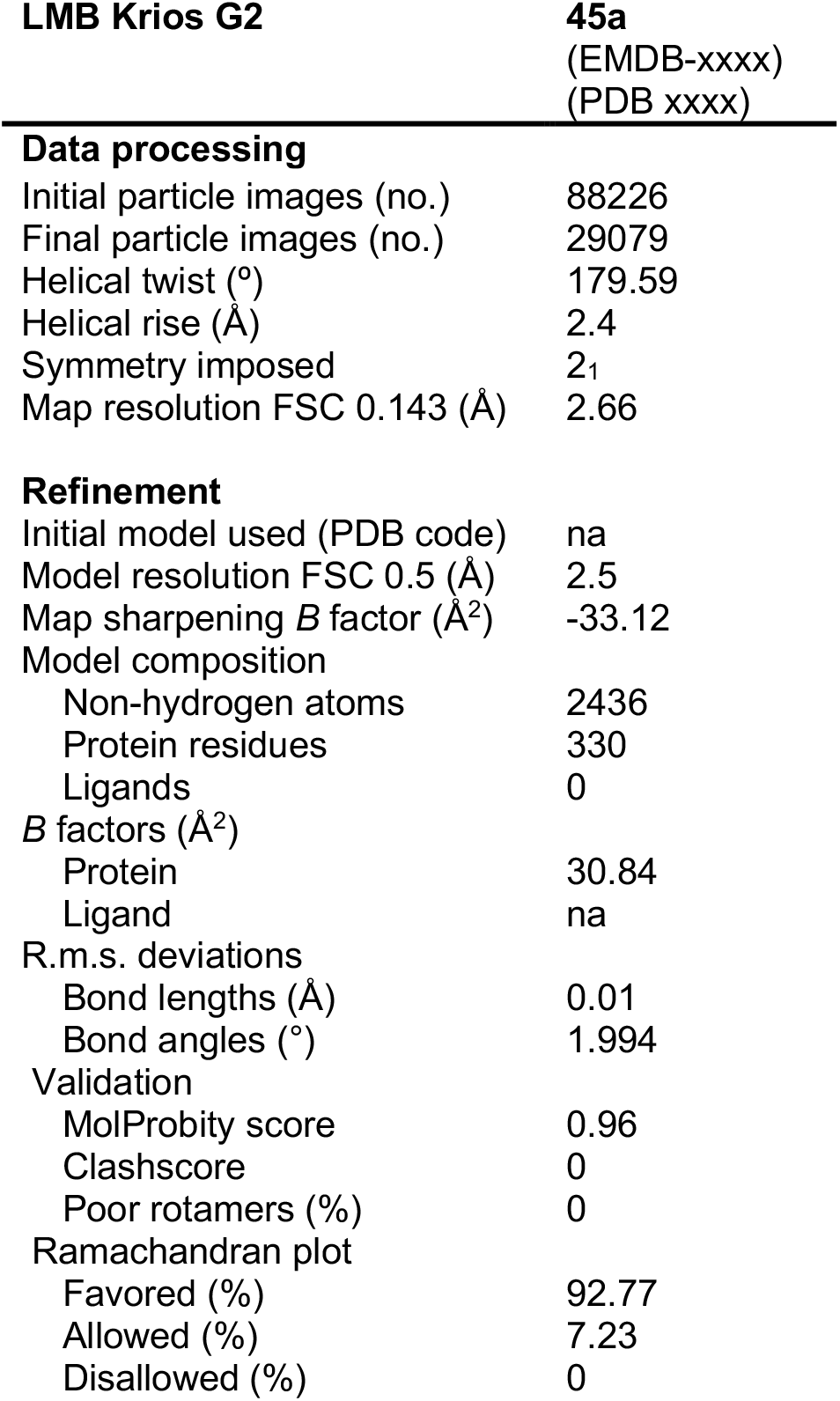

| LMB Krios G2 | 45a<br>(EMDB-xxxx)<br>(PDB xxxx) |
| --- | --- |
| **Data processing** |  |
| Initial particle images (no.) | 88226 |
| Final particle images (no.) | 29079 |
| Helical twist (°) | 179.59 |
| Helical rise (Å) | 2.4 |
| Symmetry imposed | 2<sub>1</sub> |
| Map resolution FSC 0.143 (Å) | 2.66 |
| **Refinement** |  |
| Initial model used (PDB code) | na |
| Model resolution FSC 0.5 (Å) | 2.5 |
| Map sharpening *B* factor (Å<sup>2</sup>) | -33.12 |
| Model composition |  |
| Non-hydrogen atoms | 2436 |
| Protein residues | 330 |
| Ligands | 0 |
| *B* factors (Å<sup>2</sup>) |  |
| Protein | 30.84 |
| Ligand | na |
| R.m.s. deviations |  |
| Bond lengths (Å) | 0.01 |
| Bond angles (°) | 1.994 |
| Validation |  |
| MolProbity score | 0.96 |
| Clashscore | 0 |
| Poor rotamers (%) | 0 |
| Ramachandran plot |  |
| Favored (%) | 92.77 |
| Allowed (%) | 7.23 |
| Disallowed (%) | 0 |

**Supplementary table 24.**

| TFS Krios G4 | 47a<br>(EMDB-xxxx)<br>(PDB xxxx) |
| --- | --- |
| **Data processing** |  |
| Initial particle images (no.) | 11158705 |
| Final particle images (no.) | 41080 |
| Helical twist (°) | 179.597 |
| Helical rise (Å) | 2.368 |
| Symmetry imposed | 2<sub>1</sub> |
| Map resolution FSC 0.143 (Å) | 1.86 |
| **Refinement** |  |
| Initial model used (PDB code) | na |
| Model resolution FSC 0.5 (Å) | 2.0 |
| Map sharpening *B* factor (Å<sup>2</sup>) | -21.05 |
| Model composition |  |
| Non-hydrogen atoms | 3000 |
| Protein residues | 402 |
| Ligands | 0 |
| *B* factors (Å<sup>2</sup>) |  |
| Protein | 30.44 |
| Ligand | na |
| R.m.s. deviations |  |
| Bond lengths (Å) | 0.011 |
| Bond angles (°) | 1.967 |
| Validation |  |
| MolProbity score | 1.02 |
| Clashscore | 0.34 |
| Poor rotamers (%) | 0.88 |
| Ramachandran plot |  |
| Favored (%) | 94.1 |
| Allowed (%) | 5.9 |
| Disallowed (%) | 0 |

